# Multiomic profiling of LKB1 loss of expression in breast cancer

**DOI:** 10.64898/2026.06.17.732963

**Authors:** Jinhong Kim, Ryan Holloway, Paola A. Marignani

## Abstract

The tumour suppressor LKB1 (STK11) is implicated in diverse cancers, yet its transcriptomic role in breast cancer remains poorly defined. Here, we integrate bulk-tumour genomic analysis of the METABRIC cohort, CRISPR-Cas9-mediated *STK11* knockout in human breast cancer cell lines, single-cell RNA sequencing of patient tumours, and a novel *Lkb1* murine model to characterise LKB1-dependent transcriptomic programmes across breast cancer subtypes. We discovered that the loss of STK11 induced divergent, subtype-specific gene expression changes, suppressing estrogen, progesterone and androgen signalling pathways. In patient tumours, transcriptomic intratumour heterogeneity was highest in epithelial cells, where *STK11* co-expression genes showed cell-type-selective patterns that was most striking in triple-negative breast cancer (TNBC), where STK11 was paradoxically upregulated in myoepithelial cells. While in a novel murine model, mammary-specific Lkb1 deletion drove tumourigenesis with long latency, confirming LKB1 loss as sufficient for malignant transformation with an underlying TNBC phenotype. *STK11* mutations in METABRIC samples disproportionately affected the catalytic domain in TNBC tumours and were associated with immune evasion. Together these findings highlight that loss of Lkb1 is sufficient to drive breast tumourigenesis and uncover a new role for LKB1 in TNBC.

## Introduction

Breast cancer (BC) is the most common cancer amongst women. Despite improved therapies over the last twenty years, in 2022 there were approximately 670 000 breast cancer-related deaths reported worldwide^1^. The tumour suppressor, serine-threonine kinase LKB1, also referred to as *STK11*, is associated with breast cancer^2^. Previously, we discovered that LKB1 functions as a coactivator of estrogen receptor alpha (ERα) through direct binding with the hormone receptor in the cell nucleus where it is recruited to the promoter of ERα-responsive genes. When bound to ERα, LKB1 catalytic activity enhances ERα-mediated transactivation compared with LKB1 catalytically deficient mutants^3^. More recently, we discovered the expression of the LKB1 was reduced in 31% of human epidermal growth factor receptor 2 positive (HER2+) BC and that re-engineered loss of *Lkb1* expression in an ErbB2 mouse model (*Lkb1*^−/−^/NIC) reduced the latency of mammary gland tumourigenesis where tumours were characterized as highly metabolic with elevated mTOR activity and mitochondria function^4^. Pre-clinical treatment of *Lkb1*^−/−^/NIC mice with AZD8055, that simultaneously targets mTORC1 and mTORC2, and 2-DG that targets hexokinase, significantly reduced mammary gland tumourigenesis. Simultaneous inhibition of these pathways with AZD8055/2-DG combination was significantly more effective at reducing tumour volume and burden, and activation of pro-survival pathways^5^. Previous work by others has shown that conditional *Lkb1^fl/fl^* mice^6^, crossed with ovine beta-lactoglobulin gene (BLG)-Cre mice to excise *lkb1* from mammary glands of multiparous mice, resulted in mammary gland tumourigenesis on average by 16 months^7^. In agreement with our previous findings, Klefström and colleagues analysed the role of LKB1 and c-Myc in mammary gland development and tumourigenesis with a specific emphasis on the maintenance of epithelial integrity and how LKB1 catalytic deficient mutants gain oncogenic properties, driving the expression of oncogenes^8, 9^. From these studies, functional links between LKB1 and breast health exist, thus broadening the scientific scope of LKB1 and laying the groundwork for further investigations into the role of LKB1 in breast biology. In our current study, we evaluated whether *STK11* expression is associated with mammary gland tumourigenesis at single- and bulk-cell resolutions using multi-omics approaches and confirmed that loss of *STK11* expression is sufficient to drive mammary gland tumourigenesis in a murine model.

## Methods

### *STK11* CRISPR-Cas9 cell lines

BT474 (HTB-20) and MDA-MD-231 (CRM-HTB-26) cells were purchased from ATCC and maintained according to ATCC guidelines. To generate BT474-LKB1 knockout (BT474_LKO) and MDA-MB-231-LKB1 knockout (MDA-MB-231_LKO) cells (*STK11*-knockout cells), we applied CRISPR-Cas9 technology. Single-guide RNAs (sgRNAs) targeting human *STK11* was inserted into CRISPR-Cas9 plasmid eSpCas9(1.1). No_FLAG_ATP1A1_G2_Dual_sgRNA (Addgene Plasmid #86612)-containing cells were generated. Genomic regions of interest were screened using CHOPCHOP (https://chopchop.cbu.uib.no/#) for protospacer adjacent motifs (PAMs) and sgRNA sequences. The *STK11* sgRNA (5’-CACCGTTGCGAAGGATCCCCAACG −3’) targeted the region 1207165 to 1207184 of human chromosome 19 (GRCh38.p13). Cloning protocols are described on the Addgene web site; Zhang Lab CRISPR Plasmids - Zhang Lab general cloning information (https://www.addgene.org/crispr/zhang/). Insertion of sgRNA into the eSpCas9(1.1) plasmid was confirmed by Sanger sequencing that was performed by ACGT Corporation (Toronto, Canada). Cell culture and transfection BT474 and MDA-MB-231 cells were maintained in DMEM supplemented with 8% fetal bovine serum (Invitrogen). Cell cultures were maintained at 37°C in a humidified atmosphere of 5% CO_2_. Ouabain octahydrate (Sigma) was dissolved in 95°C water at a 50mg/ml concentration and stored at –20°C. Working dilutions were prepared in water and added directly to the culture medium. Cells were electroporated using the Neon Transfection system (Invitrogen) with expression plasmids as per manufacturer’ recommendations. BT474 cells were electroporated using two pulses at 1300V for 30 milliseconds. MDA-MB-231 cells were electroporated using two pulses at 1400V for 30 milliseconds. Cells were treated with ouabain octahydrate (1µM) starting at two days post-nucleofection until all non-resistant cells were eliminated. Surveyor® assay (Integrated DNA Technologies) was carried out according to the manufacturer’s instructions using the following primers; forward primer: ACTCCACCGAGGTCATCTACC chr19:1207002-1207023 and reverse primer: AGGAAGACAGAACCATCAGCAC chr19:1207249-1207271.

### scRNA-seq from *STK11*-knockout cells

We captured single cells from each of the four breast cancer cell lines, BT474, BT474_LKO, MDA-MB-231 and MDA-MB-231_LKO, using 10x Genomics Chromium system. Following reverse transcription reaction, single-strand cDNA molecules were pre-amplified in T100 thermocycler (BioRad). To prepare cDNA libraries for next-generation sequencing (NGS) in Illumina NovaSeq 6000 system (McGill Genome Centre, QC, Canada), we followed the 10x Genomics Chromium Next Gem Single Cell 3’ Reagent Kits v3.1 (Dual Index) user guide. Raw scRNA-seq NGS reads were processed using 10x Genomics Cell Ranger v7.1, and the scRNA-seq NGS data processed were analyzed with Scanpy^10^ for differential expression analysis (DEA). In brief, following basic processing and normalization procedures, Principal Component Analysis (PCA) and Uniform Manifold Approximation and Projection (UMAP) methods were applied for dimension reduction, followed by clustering analysis using Leiden algorithm. Differentially expressed genes (DEGs) were detected by Wilcoxon test in *STK11*-knockout cells, compared with parent cells. To visualize the number of DEGs overlapping or specific to up or downregulated gene sets, we used a Venn Diagram tool (https://bioinformatics.psb.ugent.be/webtools/Venn/). To identify genes that co-expressed with *STK11* (hereafter, *STK11*-coexpression genes), we employed SciPy and collected Pearson correlation coefficient values (*r*) at the cutoff of 0.05 Bonferroni-corrected p-value following the application of Markov Affinity-based Graph Imputation of Cells (MASIC) algorithm. For Gene Set Enrichment Analyses (GSEAs), we used four programs, g:Profiler^11^, decoupler (python version)^12^, ExpressAnalyst^13^, and Metascape^14^. The g:Profiler was employed for a GSEA with a single biological database, ‘Gene Ontology Resource’ (GO), and the GSEA was performed on four DEG groups (gene sets) comprising up or downregulated genes in the two *STK11*-knockout cell lines against their parent cells. We also applied these four gene sets for a GSEA with Metascape that is comprised of multiple resource databases such GO, Kyoto Encyclopedia of Genes and Genomes (KEGG)^15^, CORUM (mammalian protein resource)^16^, WikiPathways^17^, Reactome Pathway^18^, Molecular Signatures Database transcription factor targets (MSigDB TFT)^19^, and TRRUST transcriptional regulatory networks^20^. For a GSEA with decoupler, we imported an entire read-count matrix as input data and used two reference data, 50 Hallmark gene sets (4,380 genes) and 3,061 cellular pathway gene sets (12,788 genes) available in MSigDB^21^ applying multivariate linear model (MLM). For a GSEA with ExpressAnalyst, we used Ractome Pathway database applying the four gene sets as input. In addition, proteins that are encoded by *STK11*-coexpression genes and associated with enriched Reactome pathways were linked to protein-protein interaction network (PPIN) using NDEx^22^ and Cytoscape^23^. A raw PPIN was generated using the NDEx and then processed for visualization in Cytoscape with the plugin option of yFiles Radial Layout. Codes used for bioinformatic analyses in the current study will be provided upon a request, and the processed scRNA-seq data will be uploaded to NCBI GEO database once after the current manuscript is accepted.

### Transcriptome-based intratumour heterogeneity measurement and DEA for breast cancer patients

We obtained 10x Genomics scRNA-seq data from breast cancer patients (Gene Expression Omnibus ID: GSE161529)^24^ and imported the data to Scanpy for basic processing and normalization followed by clustering analysis in four different breast cancers, 1) estrogen receptor positive (ER+), 2) HER2+, 3) *BRCA1* wild-type triple negative triple negative (TN) and 4) *BRCA1*-mutated TN (TN-BR), while normal mammary samples (normal mammoplasty) served as controls. We used decoupler for cell-type annotation on clustered cells, and detected DEGs in tumour cells compared with normal cells for each cell type using Scanpy. We selected the top 100 genes (significantly upregulated against normal) as input for a transcriptome-based intratumour heterogeneity (tITH) analysis. Gene expression profile was prepared by averaging unnormalized expression values of the selected 100 genes for individual cell types of each breast cancer. We also prepared a network file that comprises 10,892 genes and 377,669 nodes using STRING^25^ and Cytoscape^23^. The gene expression profile and network file prepared were used as input to measure tITH using nJSD^26^ in individual cell types of each breast cancer. Following the tITH measurement, we identified *STK11*-coexpression genes in each of 10 different cell types from normal mammary samples using SciPy and the MASIC algorithm and then performed a correlation analysis on each cell type of the four breast cancers using Scanpy. In addition, we selected *STK11*-involved pathway gene sets in MSigDB to use as a reference for a GSEA with decoupler. We present results from the GSEA and PPIN analysis using ExpressAnalyst and NetworkAnalyst, respectively. For a DEA to find candidate biomarker genes at bulk-cell resolution in breast cancer, we obtained mRNA expression data (data_mrna_illumina_microarray.txt) and metadata (data_mrna_illumina_microarray_zscores_ref_diploid_samples.txt) from the Molecular Taxonomy of Breast Cancer International Consortium (METABRIC) samples which are publicly available in the cBioPortal^27^ and used them to detect DEGs and find candidate biomarker genes. We performed a DEA on four different breast cancers that were grouped by immunohistochemistry (IHC)-based ER, PR, HER2 and epidermal growth factor receptor (EGFR) expression profiles referring to the ‘data_clinical_sample.txt’ file, and *STK11* mutation-containing datasets were prepared using the ‘data_mutations.txt’ file of the METABRIC sample data.

### Mice

All animal husbandry was conducted in accordance with the Canadian Council on Animal Care. Protocol #12-091 was approved by the Committee on Laboratory Animals, Dalhousie University. Hypomorphic *Lkb1^fl/fl^* ^6^, and whey acidic protein (WAP)-Cre (B6.Cg-Tg (Wap-Cre)11738Mam)^28^ were from the NCI Mouse Repository. *Lkb1^fl/fl^* mice were interbred with WAP-Cre mice to generate mice. For expression of WAP-Cre to take place, lactation is required, thus female *Lkb1^fl/fl^* /WAP-Cre were separated into two groups, the first group were nulliparous throughout the duration of the experiment, while the second group were mated for two cycles to allow for multiple pregnancies and lactations. Nulliparous female *Lkb1^fl.fl^*/WAP-Cre mice were maintained in separate cages from multiparous *Lkb1^fl/fl^*/WAP-Cre female mice. Mice were palpated every three days to monitor for mammary tumours, grooming abnormalities and weight change.

### Whole Mounts and Histology

Inguinal mammary glands were removed and mounted onto glass slides, then fixed in Carnoy’s solution overnight. Following this, glands were hydrated and stained with Carmin alum, dehydrated and mounted as described^29^. Mammary gland were prepared for IHC analysis previously described^4^. Formalin fixed and paraffin embedded tissues were prepared for histology followed by hematoxylin and eosin (H/E) staining. Tissue sections were prepared and imaged as previously described^3–5^. Tissue sections were incubated in primary antibodies: LKB1 (Santa Cruz) at a dilution of 1:30, cyclin D, c-Myc, HIF1, LDH, pS6 (S235/S236), ErbB2, PGR and ERα at a dilution of 1:100 (Cell Signaling). Images were obtained using Nikon Eclipse TE 2000-E, mounted with a Q-Imaging CCD camera and acquired using the Simple PCI software as previously described^3^.

### Statistical Analysis

Results were derived from a minimum of three independent experiments. For Kaplan-Meir curves, Log ranked tests, scores of 3 or higher were considered significant (p<0.0001). Tissues were harvested from three separate C57BL/6J wild-type mice, three separate multiparous *Lkb1^fl/fl^*/WAP-Cre and three separate LKB1*^fl/fl^* mice. To screen significant DEGs and/or GSEA outputs generated from scRNA-seq or microarray data, we applied the cutoff of 0.05 p-value corrected (or adjusted) for false discovery rate (FDR) using Benjamini-Hochberg or Bonferroni method.

## Results

### scRNAseq analysis of *STK11*-knockout breast cancer cells

To characterize how the loss of LKB1 changes transcriptomic profiles of human breast cancers, we created *STK11*-knockout cells from two genetically distinct breast cancer cell lines, BT474 (ER+/PR+/HER2+) and MDA-MB-231 (ER^−^/PR^−^/HER^−^) using CRISPR-Cas9 technology. Western blot analysis confirmed complete knockout of LKB1 (Fig. 1a) from the two cell lines resulting in loss of *STK11* expression in BT474 (BT474_LKO; −1.36 log_2_ fold change) and MDA-MB-231 (MDA-MB_231_LKO; −1.98 log_2_ fold change), compared with parent cells (Source Data Fig. 1). For differential expression analysis (DEA) in whole transcriptomes of individual single cells, we obtained >1.53 billion raw scRNA-next generation sequencing (NGS) reads from 17,575 single cells from the four cell lines (Supplementary Table 1) and performed clustering analysis on processed scRNA-seq NGS data (Fig. 1b). The DEA identified similar number of up- or down- regulated genes in BT474_LKO, while MDA-MB-231_LKO showed 1.7 times more downregulated genes than upregulated genes when compared with corresponding parent cells indicating a stronger directional effect on gene expression in TNBC following the *STK11* knockout (Fig. 2c). Heatmap analysis presents normalized expression values (z-scores) of the most highly expressed top 50 genes in each of the four cell lines, showing overall similarity in the extent of gene expression values between parent and *STK11*-knockout cells and confirmed the genetic difference between BT474 and MDA-MB-231 cell lines (Fig. 1d). These distinct profiles in gene expression between BT474 and MDA-MB-231 also affected identification of few overlapping differentially expressed genes (DEGs) (Extended Data Fig. 1a). Based on the extent of log_2_ fold change values of DEGs, heatmap analysis presents top 10 up- or down- regulated genes in each of the four cell lines (Extended Data Fig. 1b), and an expression pattern for 5 of the top 10 genes, where expression was cell line specific, are presented in single cells within UMAP plots (Extended Data Fig. 1c,d). When we compared a pattern of gene expression for two hormone receptors (HRs; *ESR1* and *PGR*) and two receptor tyrosine kinases (RTKs; *ERBB2* and *EGFR*) in parent versus *STK11*-knockout cells, we observed enhanced expression of *PGR* (0.38 log_2_ fold change) and *ERBB2* (0.43 log_2_ fold change) and decreased expression of *EGFR* (−1.20 log_2_ fold change) in response to loss of LKB1 in BT474. In contrast, only *EGFR* expression (0.39 log_2_ fold change) increased in MDA-MB-231_LKO (Fig. 1e and Source Data Fig. 1). These results suggest that the loss of LKB1 leads to a change in expression of HR and RTK genes in breast cancer epithelial cells. Furthermore, a GSEA using Metascape with multiple databases of genome annotation on the four cell lines identified functional gene sets associated with loss of LKB1 (see ‘Methods’ for more detailed information). Interestingly, we found common downregulation of cell cycle-related biological processes in both *STK11*-knockout cell lines (Fig. 1f), and the GSEA using two different transcription factor databases confirmed the downregulation of cell cycle (MSigDB TFT ID: M30131) and DNA replication (TRRUST ID: TRR00781) (Extended Fig. 1e and Source Data Extended Fig. 1). When cellular adenosine monophosphate (AMP) concentration increases, LKB1-STRAD-MO25 complex phosphorylates AMP-binding AMPK, thereby regulating cellular adenosine triphosphate (ATP) concentration for cell proliferation and cycling processes, and these generate ATP molecules through glycolysis or fatty acid oxidation^30^. The loss of LKB1, however, induces metabolic stress condition by accelerating ATP consumption and consequently suppresses cell cycle process under ATP depletion because of dysfunctional ATP supplement processes such as no phosphorylation of unc-51 like autophagy activating kinase 1 (ULK1) by LKB1 in autophagy, active acetyl-coA carboxylase not inhibited by LKB1-phosphorylated AMPK in fatty acid oxidation and decreased aerobic glycolysis (Warburg effect) (Table 1 and Source Data Fig. 1). In addition, LKB1 is required to maintain mitotic spindle orientation ^31^ and we identified that the loss of LKB1 led to downregulation of centrosome separation (Gene Ontology ID: GO:0051294), microtubule cytoskeleton organization (GO:0000226), negative regulation of progression through cell cycle (GO:0045786) and regulation of cyclin-dependent protein kinase activity (GO:0008064). These profiles confirm that the loss of LKB1 negatively affects cell cycle as well as stable cell division processes (Source Data Fig. 1).

**Fig. 1:**
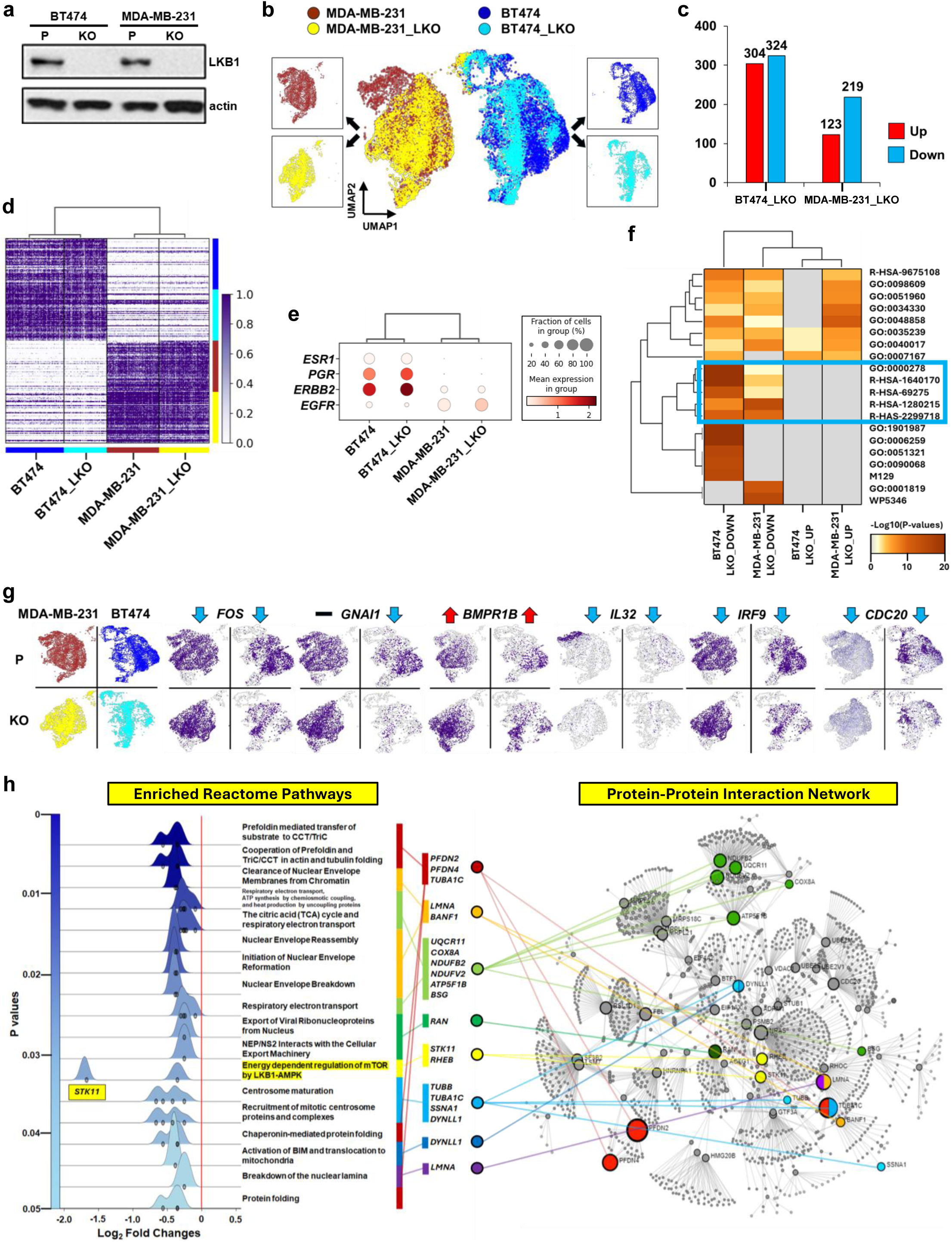
Profiles of differential gene expression at single-cell resolution in *STK11*-knockout breast cancer. **a,** Western blotting analysis. Absence of *STK11-*encoded protein (LKB1) expression was confirmed in BT474 and MDA-MB-231 breast cancer cell lines. **b**, Single-cell clusters of *STK11*-knockout and wild type breast cancer cells. UMAP plot presents clusters of single cells from BT474 parent (blue), BT474_LKO (cyan), MDA-MB-231 parent (brown) and MDA-MB-231_LKO (yellow) cell lines. Black arrows point to cells for each cell line. **c,** X and y axes indicate *STK11*-knockout BT474 and MDA-MB-231 breast cancer cell lines and the number of up (red) and down (blue) regulated genes in BT474_LKO and MDA-MB-231_LKO, respectively. **d**, Gene expression in the four breast cancer cell lines. Heatmap presents normalized expression values of top 50 significantly upregulated genes per cell line. Two colour-matching bars at right and bottom indicate each of the four cell lines. A white-purple colour bar shows low and high z-scores of normalized gene expression values, respectively. **e,** Gene expression profiles of two hormone receptors and two receptor tyrosine kinases. The extent of normalized gene expression values for these four genes is presented on dot plot (*ESR1*: estrogen receptor 1, *PGR*: progesterone receptor, *ERBB2*: erbb2 receptor tyrosine kinase 2, and *EGFR*: epidermal growth factor receptor, respectively). A white-dark brown colour bar indicates low (0), medium (1) and high (2) normalized mean expression values, and a gradient size of grey dots represents the fraction (%) of cells where individual genes are expressed. **f**, Top 20 enriched gene sets from gene set enrichment analysis (GSEA) with up- or down-regulated genes at ≥ |0.58| log_2_ fold change (≈ 1.5 times fold change) in BT474_LKO and MDA-MB-231_LKO. A grey-brown colour bar shows statistical significance with negative log_10_(p-values). Blue square line highlights gene ontology biological processes or pathways commonly downregulated in the two *STK11*-knockout cell lines (R-HSA-9675108: Nervous system development, GO:0098609: Cell-cell adhesion, GO:0051960: Regulation of nervous system development, GO:0034330: Cell junction organization, GO:0048858: Cell projection morphogenesis, GO:0035239: Tube morphogenesis, GO:0040017: Positive regulation of locomotion, GO:0007167: Enzyme-linked receptor protein signalling pathway, GO:0000278: Mitotic cell cycle, R-HSA-1640170: Cell cycle, R-HSA-69275: G2/M transition, R-HSA-1280215: Cytokine signalling in immune system, R-HAS-2299718: Condensation of prophase chromosomes, GO:1901987: Regulation of cell cycle phase transition, GO:0006259: DNA metabolic process, GO:0051321: Meiotic cell cycle, GO:0090068: Positive regulation of cell cycle process, M129: PID PLK1 pathway, GO:0001819: positive regulation of cytokine production, and WP5346: 8p23 1 copy number variation syndrome). See the Source Data Extended Data Fig. 1 for detailed GSEA information. **g**, Profiles of representative DEGs. Purple colour dots on UMAP plots indicate single cells where genes are expressed in parent cell lines (P) or *STK11*-knockout cells (KO) from MDA-MB-231 (left) and BT474 (right). Black dash presents non-significant change, while blue and red arrows represent down and upregulation of individual genes, respectively (*FOS*: Fos proto-oncogene, *GNAI1*: G protein subunit alpha i1, *BMPR1B*: bone morphogenetic protein receptor type 1B, *IL32*: interleukin 32, *IRF9*: interferon regulatory factor 9, and *CDC20*: cell division cycle 20). **h**, Enriched cellular pathways and protein-protein interaction network. A blue-gradient bar on *y* axis of ridgeline plot (left) shows p-values (< 0.05) for Reactome pathways enriched with *STK11*-coexpression genes. X axis presents averaged log_2_ fold changes of *STK11*-coexpression genes in *STK11*-knockout cells when compared with parent cells of BT474 and MDA-MB-231 cell lines. A list of enriched pathways and associated genes are presented adjacent to the ridgeline plot. Multiple colour bars indicate a group of genes in individual enriched pathways. Colour-matching lines connect individual gene groups to location of their encoded proteins in a protein-protein interaction network (right) (*PFDN2*: prefoldin subunit 2, *PFDN4*: prefoldin subunit 4, *TUBA1C*: tubulin alpha 1c, *LMNA*: lamin A/C, *BANF1*: barrier to autointegration nuclear assembly factor 1, *UQCR11*: ubiquinol-cytochrome c reductase, complex III subunit XI, *COX8A*: cytochrome c oxidase subunit 8A, *NDUFB2*: NADH:ubiquinone oxidoreductase subunit B, *NDUFV2*: NADH:ubiquinone oxidoreductase core subunit V2, *ATP5F1B*: ATP synthase F1 subunit beta *BSG*: basigin (Ok blood group), *RAN*: RAN, member RAS oncogene family, *STK11*: serine/threonine kinase 11, *RHEB*: Ras homolog, mTORC1 binding, *TUBB*: tubulin beta class I, *SSNA1*: SS nuclear autoantigen 1, and *DYNLL1*: dynein light chain LC8-type 1).

**Fig. 2:**
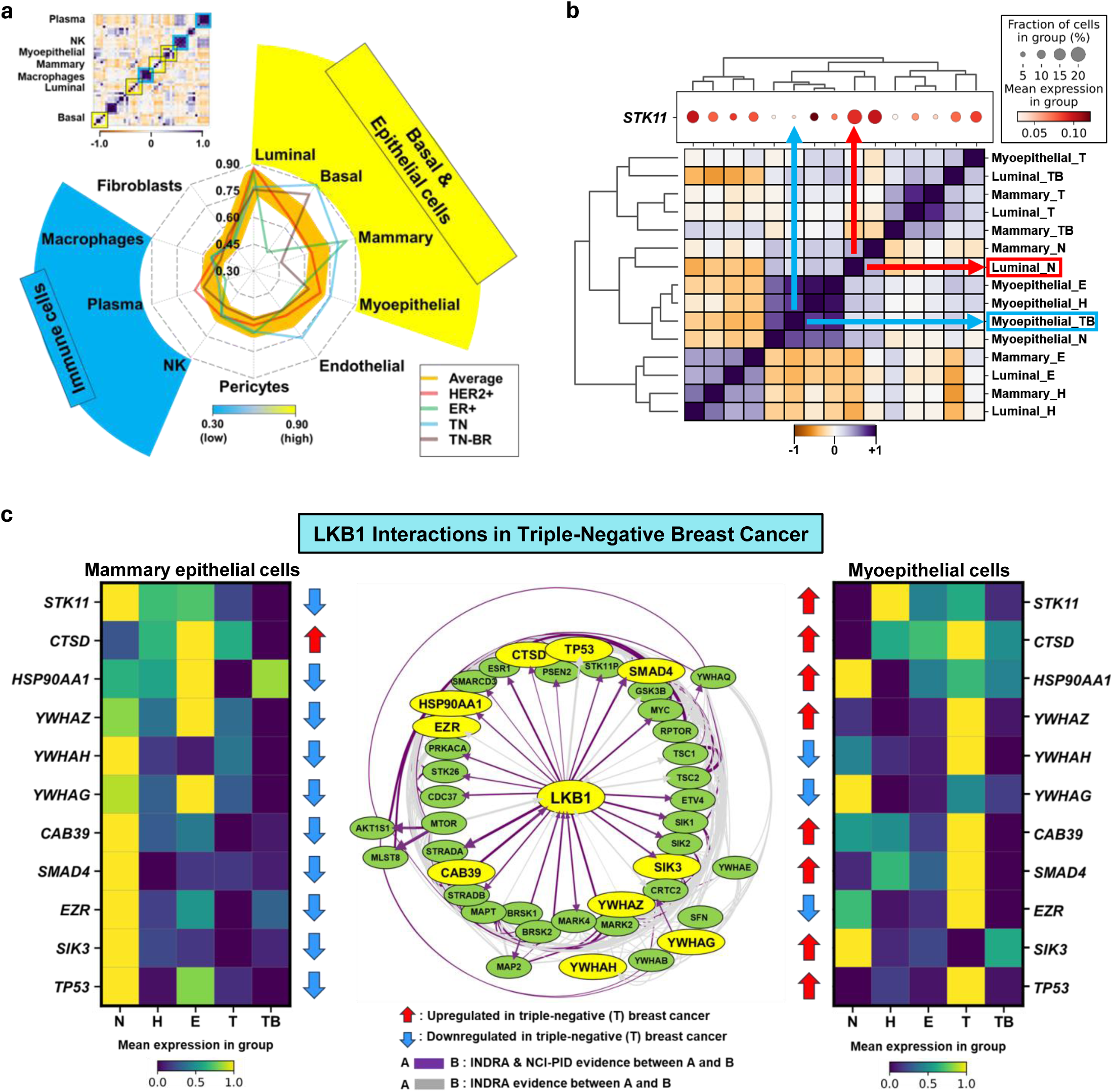
Profiles of differential gene expression at single-cell resolution in various cell types of breast cancers. **a**, Transcriptome-based intratumour heterogeneity (tITH) in breast cancers. The numbers in the centre of net plot indicate the extent of tITH in 10 different cell types of HER2+ (red), ER+ (green), *BRCA1*-wild type triple negative (TN, blue), and *BRCA1*-mutated triple negative (TB-BR, brown) breast cancers. A wide-orange line indicates the averaged tITH of the four breast cancers. A blue-yellow colour bar represents low (0.30) and high (0.90) tITH, respectively. The colour-matching blue and yellow sections highlight cell types that have relatively low and high tITH, respectively. Blue and yellow colour-matching squares in the correlation-matrix plot located at the left top indicate hierarchically clustered cell types from the four breast cancers. An orange-white-purple colour bar represents low (−1.0), medium (0) and high (1.0) z-scaled correlation scores, respectively (NK: natural killer cells). **b**, Gene expression profiles in breast epithelial cells. A white-dark brown colour bar indicates low (0.05) and high (0.10) normalized mean expression of *STK11*. A gradient size of grey dots represents the fraction (%) of cells where *STK11* was expressed in three types of breast epithelial cells (luminal, mammary and myoepithelial) from the normal mammary (N) and four different breast cancers (E: ER+, H: HER2+, T: *BRCA1* wild-type triple negative, and TB: *BRCA1*-mutated triple negative). The order of cell type-sample list from top to bottom in the right of correlation matrix is symmetric to red-dot plot from left to right. An orange-white-purple colour bar represents low (−1.0), medium (0) and high (+1.0) z-scaled correlation scores of S*TK11*-coexpression genes. The red and blue arrows and squares indicate the highest and lowest fractions of epithelial cells, respectively, where *STK11* is expressed. **c**, Profiles of *STK11*-coexpression genes associated with LKB1 pathway. Two purple-green-yellow colour bars indicate low (0), medium (0.5) and high (1.0) z-scaled mean expression values of 11 differentially expressed and *STK11*-coexpression genes, including *STK11*, that are associated with the LKB1 pathway in *BRCA1* wild-type triple-negative breast cancer (T). The matrix plots present differential expression of the genes in mammary epithelial (left) and myoepithelial (right) cells, for each of the five different breast samples. The red and blue arrows indicate up and downregulation of the genes, respectively, in the triple-negative breast cancer when compared with normal mammary cells. Purple and grey lines in the protein-protein interaction network (PPIN; Centre) represent if the interaction evidence is based on both ‘Integrated Network and Dynamical Reasoning Assembler (INDRA)’ and ‘National Cancer Institute Pathway Interaction Database (NCI-PID)’ systems (purple) or only INDRA system (grey). Yellow and green oval-shape nodes in the PPIN present the profiled 11 genes and rest of genes in the LKB1 pathway, respectively (*STK11*: serine/threonine kinase 11, *CTSD*: cathepsin D, *HSP90AA1*: heat shock protein 90 alpha family class A member 1, *YWHAZ*: tyrosine 3-monooxygenase/tryptophan 5-monooxygenase activation protein zeta, *YWHAH*: tyrosine 3-monooxygenase/tryptophan 5-monooxygenase activation protein eta, *YWHAG*: tyrosine 3-monooxygenase/tryptophan 5-monooxygenase activation protein gamma, *CAB39*: Calcium Binding Protein 39, *SMAD4*: SMAD family member 4, *EZR*: ezrin, *SIK3*: SIK family kinase 3, *TP53*: tumour protein p53).

**Table 1.** Hallmark gene sets enriched in breast cancer. Gene set enrichment analysis (GSEA) was performed on *STK11*-knockout breast cancer cell lines (BT474_LKO and MDA-MB-231_LKO) comparing with parent cells using 50 Hallmark gene sets available in MSigDB. The table contains the most significantly downregulated 10 hallmark gene sets. Notably, five gene sets are identical between the two *STK11*-knockout cell lines when sorting gene sets from the smallest log_2_ fold change values. A full list of the Hallmark gene sets enriched is available in Source Data Fig. 2.

We selected representative DEG markers associated with functional pathways enriched through a gene set enrichment analysis (GSEA) using canonical pathway gene sets available in MSigDB database. The first marker is *FOS* that encodes for the protein, fos proto-oncogene, AP-1 transcription factor subunit, negatively regulating *PGR* expression in breast cancer^32^. *STK11* knockout downregulated *FOS* three times lower in BT474_LKO (−1.24 log_2_ fold change) than MDA-MB-231_LKO (−0.36 log_2_ fold change), and *PGR* was upregulated (0.38 log_2_ fold change) in BT474_LKO (Fig. 1g and Source Data Fig. 1) with non-significant change in MDA-MB-231_LKO. The inverse expression pattern between *FOS* and *PGR* confirms the reported biological function of *FOS* in the estrogen-induced *PGR* expression regulation. The GSEA also revealed that *STK11* knockout suppressed the estrogen-dependent gene expression pathway (MSigDB ID: R-HSA-9018519; −0.65 and −0.53 log_2_ fold changes in BT474_LKO and MDA-MB-231_LKO, respectively) and extra nuclear estrogen signalling pathways (MSigDB ID: R-HSA-9009391; −0.36 and −0.24 log_2_ fold changes in BT474_LKO and MDA-MB-231_LKO, respectively) (Source Data Fig. 1). *FOS* was one of the most significantly downregulated genes in the two estrogen-related pathways (Source Data Fig. 1), while *ESR1* showed no significant change in *STK11*-knockout cells of BT474 and MDA-MB-231. These gene profiles collectively suggest that LKB1 is positively correlated with FOS in regulating progesterone-medicated signalling pathways in breast cancers. The second marker is *GNAI1* that was the most significantly downregulated gene (−1.96 log_2_ fold change in BT474_LKO) in the extra nuclear estrogen signalling pathway, followed by *FOS* (Fig. 1g and Source Data Fig. 1). *GNAI1* encodes for the protein, G protein subunit alpha i1, that is involved in G protein-coupled estrogen signalling pathways through plasma membrane^33^. These profiling results support our previous findings that show LKB1 is involved in non-genomic estrogen-mediated signalling events in addition to coactivating ERα in cell nucleus^3^. *BMPR1B* encoding for the protein, bone morphogenetic protein receptor type 1B, an androgen-responsive gene^34^, and found to be upregulated in *STK11*-knockout cells, with higher expression in BT474_LKO (1.42 log_2_ fold change) compared to MDA-MB-231_LKO (0.77 log_2_ fold change) (Fig. 1g and Source Data Fig. 1). These finding suggest that LKB1 expression is negatively correlated with BMP receptor expression in breast cancers as in lung cancer. SMAD protein family members are phosphorylated by BMPR1B, and when phosphorylated, binds to androgen receptors (ARs) where the receptors are functionally antagonized^35^. Although *AR* gene expression itself showed an opposite pattern, that is, downregulation (−0.57 log_2_ fold change) and upregulation (0.45 log_2_ fold change) in BT474_LKO and MDA-MB-231_LKO, respectively, *STK11* knockout supressed the gene set of AR signalling pathway (WikiPathways ID: WP138) in both BT474_LKO (−0.40 log_2_ fold change) and MDA-MB-231_LKO (−0.36 log_2_ fold change) (Source Data Fig. 1). These profiles indicate that LKB1 positively regulates androgen-medicated signalling pathways in breast cancers. *IL32* encodes for the protein, interleukin 32, that is associated with breast cancer metastasis^36^. *IL32* was downregulated in BT474_LKO and MDA-MB-231_LKO showing more than three times lower expression in MDA-MB-231_LKO (−2.80 log_2_ fold change) than BT474_LKO (−1.10 log_2_ fold change) (Fig. 1g and Source Data Fig. 1). *IL32* is involved in cytokine signalling in the immune system (Reactome Pathway ID: R-HSA-1280215) that was downregulated in both BT474_LKO and MDA-MB-231_LKO (−0.72 and −2.04 log_2_ fold changes, respectively) (Fig. 1f and Source Data Fig. 1). Although *IL32* and the cytokine signalling pathways were downregulated in both *STK11*-knockout cell lines, suppressors of cytokine signalling (SCOS) gene family showed an opposite expression pattern, that is, downregulation of *SOCS1* (−1.02 log_2_ fold change) in MDA-MB-231_LKO with no significant change in BT474_LKO and upregulation of *SOCS2* (0.88 log_2_ fold change) in BT474_LKO. *CCL2* encodes for the protein, C-C motif chemokine ligand 2, and is involved in the pathway of cytokine signalling in the immune system (Source Data Fig. 1). CCL2 is also associated with breast cancer metastasis^37^, and *CCL2* expression was inversed when comparing with *SOCSs*, that is, downregulation (−1.77 log _2_ fold change) and upregulation (2.80 log_2_ fold change) in BT474_LKO and MDA-MB-231_LKO, respectively (Source Data Fig. 1). This indicates that LKB1 is positively correlated with Interleukin 32 (IL32) and cytokine signalling pathway, but cytokine-induced immune cascades in breast cancers are differentially responded to the presence or absence of HRs or RTKs. In addition to *FOS* and *IL32*, interferon regulatory factor 9 (*IRF9*) is also involved in cytokine signalling in the immune system (Source Data Fig. 1), and was co-downregulated (−0.81 log_2_ fold change and −0.65 log_2_ fold change in BT474_LKO and MDA-MB-231_LKO, respectively) in the two *STK11*-knockout cell lines (Fig. 1g and Source Data Fig. 1). The GSEA using TRRUST database revealed that the BRCA1-regulated gene set (TRRUST ID: TRR00075), including *IRF9* and *FOS*, was downregulated in BT474_LKO with no significant change in MDA-MB-231_LKO (Source Data Extended Fig. 1). *IRF9* is required for apoptosis of malignant cells that is primed by BRCA1 collaborating with *IRF7*^38^. However, an expression pattern of *BRCA1* and *IRF7* was variable in the current study, that is, downregulation (−0.36 log_2_ fold change) of *BRCA1* in BT474_LKO with no significant change in MDA-MB-231_LKO and upregulation (0.28 log_2_ fold change) of *IRF7* in MDA-MB-231_LKO with no significant change in BT474_LKO (Source Data Fig. 1). These profiling results confirm that the loss of LKB1 is indirectly associated with immune responses in breast cancers. Of interest was Cell Division Cycle 20 (*CDC20)* that was downregulated in BT474_LKO (−1.08 log_2_ fold change) and MDA-MB-231_LKO (−0.14 log_2_ fold change) (Source Data Fig. 1). *CDC20* belongs to three MSigDB Hallmark gene sets, MYC_TARGETS_V1 (MSigDB ID: M5926), E2F_TARGETS (M5925) and G2M_CHECKPOINT (M5901), which were downregulated in the two *STK11*-knockout cell lines (Table 1 and Source Data Fig. 1). These results suggest that LKB1 is positively associated with transcriptional activities, initiation of mitosis and DNA damage-repair processes, and that the loss of LKB1 leads to dysfunctional regulation of these processes.

### *STK11*-coexpression genes

We performed a co-expression analysis on parent cells to identify biological pathway gene sets involved in breast cancer that show a similar expression pattern to *STK11* (*STK11*-coexpression genes). For downstream analysis, we selected 92 *STK11*-coexpression genes that were commonly found in BT474 and MDA-MB-231 cell lines excluding 14 ribosomal RNA genes (Source Data Fig. 2). For example, the top five representative *STK11*-coexpression genes that are significantly correlated with *STK11* expression are splicing factor 3b subunit 2 (*SF3B2*; average of imputed Pearson correlation coefficient values in two parent cells: 0.7344), high mobility group 20B (*HMG20B*; 0.7284), NRAS proto-oncogene, GTPase (*NRAS*; 0.7184), heparin binding growth factor (*HDGF*; 0.7041), and cyclase associated actin cytoskeleton regulatory protein 1 (CAP1; 0.6878) (Source Data Fig. 1). We identified that most *STK11*-coexpression genes were downregulated with the exception of eukaryotic translation initiation factor 4A2 (*EIF4A2*), a key regulator of mTOR, was upregulated (0.73 and 0.26 log_2_ fold change in BT474_LKO and MDA-MB-231_LKO, respectively) (Source Data Fig. 1). *EIF4A2* is associated with cell proliferation and chemosensitivity in TNBC^39^, and the gene expression profiles in the current study indicate that LKB1 negatively regulates EIF4A2 regardless of the presence and absence of HRs and RTKs supporting the tumour-suppressive function of LKB1 in breast cancers. For functional characterization of *STK11*-coexpression genes, we performed a GSEA on the 92 selected *STK11*-coexpression genes using Reactome pathway database. A total of 18 cellular pathways were enriched with 17 *STK11*-coexpression genes, including the pathway of ‘energy dependent regulation of mTOR by LKB1-AMPK’, where *STK11* is directly involved (Fig. 1h and Source Data Fig. 1). In addition, we performed a PPIN analysis on the 92 *STK11*-coexpression genes and found that all the 17 *STK11*-coexpression genes are functionally connected in a PPIN (Fig. 1h).

### Transcriptomic profiles at single-cell resolution in epithelial cells of breast cancer patients

In the previous section, we profiled DEGs by comparing epithelial cells of *STK11*-knockout breast cancer cell lines to those of parent cell lines and associated them to biological processes where LKB1 is directly or indirectly correlated. To validate these findings in multiple cell types, we performed a DEA on scRNA-seq data generated from a total of 28 samples comprising 8 normal mammary tissues from reduction mammoplasties of women with no family history of breast cancer and 20 primary breast tumour tissues of treatment-naïve patients (Source Data Fig. 2). First, to characterize transcriptomic complexity in breast cancers, we measured tITH using nJSD^40^ in each of 10 different cell types from ER+, HER2+, TN (*BRCA1*-wild type) and TN-BR (*BRCA1*-mutated) breast cancers. For this measurement, we stratified the most significantly upregulated top 100 genes in tumour cells compared with genes from normal mammary cells (Source Data Fig. 2). For each of the four breast cancers, expression values of the top 100 genes in each cell type were averaged to calculate tITH values. We found three epithelial cell types, luminal, mammary and myoepithelial cells, with basal cells showed relatively higher tITH (averaged values: 0.66 ∼ 0.81) than immune cells, such as natural killer (NK; 0.59), plasma (0.55) and macrophages (0.53) (Fig. 2a and Source Data Fig. 2). The highest and lowest tITHs were found in luminal epithelial cells (0.81) and fibroblasts (0.51), respectively (Source Data Fig. 2). Both ER+ and TN-BR showed relatively more variations on tITH in mammary epithelial and basal cells, when compared with other breast cancers (Source Data Fig. 2). For example, in TN-BR mammary epithelial cells, tITH was 0.46, while tITHs of the cells in ER+ was 0.85 (Source Data Fig. 2). To identify relationships between tITH values and *STK11* expression, we prepared a set of 1,864 *STK11*-coexpression genes in 10 common cell types that were also DEGs in each cell type of the four breast cancers when compared with normal mammary cells (Source Data Fig. 2). A correlation analysis with *STK11*-coexpression genes (in the left top of Fig. 2a) revealed high similarity between the extent of tITH and gene expression profiles (Fig. 2a). This represents a positive correlation between transcriptomic heterogeneity and a pattern of *STK11*-coexpression genes in the breast cancers we analysed. The tITH analysis showed a relatively higher heterogeneity (or complexity) in epithelial cells when compared with immune and other cells in transcriptomes of breast cancers. We further investigated which epithelial cell types showed more positive correlation with expression pattern of *STK11*-coexpression genes including *STK11* in normal and tumour mammary samples. We profiled *STK11* expression in luminal, mammary and myoepithelial cells and found *STK11* expression was relatively higher in the three epithelial cells from normal mammary samples, than tumour ones (Fig. 2b). Luminal epithelial cells from normal mammary samples have the largest fraction of cells (∼20%) where *STK11* was expressed, while TN-BR myoepithelial cells show the lowest cell fraction (<5%) where *STK11* was expressed. For most tumour epithelial cells, *STK11* was downregulated compared with normal mammary cells. However, *STK11* was more than 2.8 times (1.49 log_2_ fold change) upregulated in TN myoepithelial cells when compared with normal myoepithelial cells (Fig. 2b and Source Data Fig. 2). Overall, most types of TN and TN-BR epithelial cells, except TN-BR myoepithelial cells, displayed a distinct transcriptomic profile compared with epithelial cells from other breast cancers (Fig. 2b), and myoepithelial cells from most breast cancers, except from the TN breast cancer, showed higher transcriptomic similarity than other cell types (Fig. 2b).

### *STK11* expression and TNBCs

Based on the profiles of *STK11* and *STK11*-coexpression genes in breast cancer patient samples, TN shows a unique expression pattern of the genes, particularly in mammary and myoepithelial cells. For example, *STK11* was more than 2.3 times (−1.24 log_2_ fold change) and 3.6 times (−1.87 log_2_ fold change) downregulated in mammary epithelial cells of TN and TN-BR, respectively, when compared with normal mammary cells (Source Data Fig. 2). In contrast, *STK11* was 2.8 times (1.49 log_2_ fold change) upregulated in TN myoepithelial cells (Source Data Fig. 2) when compared with normal myoepithelial cells, and an expression pattern of the gene did not show a significant change in TN-BR myoepithelial cells. We performed a GSEA on *STK11*-coexpression genes using MSigDB pathway database to identify cellular pathways where *STK11*-coexpression genes are functionally associated in mammary epithelial and myoepithelial cells of TN breast cancer. Of 16 *STK11*-associated pathways, we observed a significant decrease in activity of the LKB1 pathway (PID_LKB1_Pathway) in HER2+, TN and TN-BR (−1.25, −1.19 and −1.01 log_2_ fold changes, respectively) breast cancers when compared with normal mammary cells (Source Data Fig. 2). The LKB1 pathway is comprised of 47 genes (Source Data Fig. 2), of which 15 were *STK11*-coexpression genes, including *STK11* (Source Data Fig. 2), and 45 of the 47 genes were DEGs in mammary epithelial and/or myoepithelial cells (Source Data Fig. 2). We profiled 11 genes that commonly belong to the groups of DEGs and *STK11*-coexpression genes in the two epithelial cell types of TN breast cancer. The correlation and PPIN analyses revealed that the 11 genes profiled are functionally centred in the LKB1 pathway presenting an inverse expression pattern between the mammary epithelial and myoepithelial cell types in the TN breast cancer (Fig. 2c). For example, 10 of the 11 genes, except cathepsin D (*CTSD*), were downregulated in mammary epithelial cells, while 7 of the 10 genes were upregulated in myoepithelial cells with *STK11* (Fig. 2c). These profiling results suggest that LKB1 may actively function as a tumour suppressor in myoepithelial cells of TN breast cancer.

### *STK11* mutations are associated with differential gene expression in breast cancers

Through profiling scRNA-seq data in the previous two sections, we discovered that LKB1 is associated with a variety of biological processes in breast cancers, particularly in TN breast cancer. Here, we expanded our investigation to characterize the relationships between mutational status of *STK11* and differential gene expression at bulk-tumour resolution in breast cancers. The cancer genomics database, cBioPortal^27^, has 2,509 METABRIC samples, of which 1,980 samples include normalized microarray mRNA expression data. Based on the expression pattern of IHC-confirmed ER, PR, and HER2 with information on *STK11* mutations, 1,734 of the 1,980 METABRIC samples were grouped into the following four breast cancer types; 1) 930 (49.84%) samples for ER+/PR+/HER2− (ER+/PR+), 2) 401 (21.49%) samples for ER+/PR-/HER2− (ER+), 3) 125 (6.70%) samples for ER-/PR-/HER2+ (HER2+), and 4) 278 (14.90%) samples for ER- /PR-/HER2− (*BRCA1-*WT triple negative; TN) (Source Data Fig. 3). We performed a clustering analysis on the 1,734 mRNA expression data and found that HER2+ and TN samples were separated from ER+/PR+ or ER+ samples, and PR status had less impact on transcriptomic changes, that is, most ER+/PR+ and ER+ samples overlapped (Fig. 3a). We observed a positive correlation between an expression of HR and RTK genes and sample distribution (Fig. 3b). This clustering analysis based on profiles of differential gene expression confirmed relatively higher transcriptomic similarity between ER+/PR+ and ER+ breast cancers, when compared with HER2+ or TN breast cancer at bulk-cell resolution. From a deeper analysis of the METABRIC samples, we found 21 mutations in *STK11* (Source Data Fig. 3). The *STK11* mutations were detected in three different domains; 2 mutations within the N-terminus domain (1 ∼ 51 amino acids; aa), 13 mutations within the protein kinase domain (52 ∼ 309 aa) and 6 mutations within the C-terminus domain (310 ∼ 433 aa). More specifically, *STK11* mutations comprise 13 missense, 7 truncation and 1 splice junction. To present relationships between mutational status of *STK11* and differential gene expression, we performed a correlation analysis on METABRIC samples dividing into three groups based on mutation sites; 13 samples for *STK11* mutation within the protein kinase domain, 8 samples for *STK11* mutation within the N- and C-termini non-kinase domain, and 1,959 samples for *STK11* wild-type (WT). Interestingly, we found relatively higher correlation within TN samples that comprise 6 *STK11-*kinase domain mutants and 1 *STK11*-non-kinase domain mutant with 1 *STK11*-kinase domain mutant from ER+ sample (Fig. 3c). Corresponding to a distinct distribution of TN METABRIC samples shown in the UMAP plot (Fig. 3a), the correlation analysis results support an overall transcriptomic uniqueness in TN breast cancer when compared with other breast cancers. These data also highlight a clear difference in a pattern of gene expression between *STK11*-mutated and *STK11-*WT breast cancers, with a particular emphasis on unique profile of gene expression in TN breast cancer. For example, TN breast cancer has a relatively higher portion (6; 85.71%) of *STK11* mutations within the protein-kinase domain compared with all other mutations identified in the METARIC dataset (1 mutation from HER2+, 2 mutations from ER+/PR+, and 3 mutations from ER+ breast cancers) (Source Data Fig. 3). Of the 21 *STK11* mutations, 13 and 2 mutations were identified in early- (Stage 1 and 2) and late- stage (Stage 3) cancers, respectively, and the remaining 6 mutations were identified at unknown staging of breast cancers. For more specific DEA related to mutational status of *STK11* and cancer stage, we profiled DEGs of early-stage METABRIC samples that contain the 13 *STK11* mutations in ER+/PR+, ER+ and TN breast cancers. We compared normalized mRNA expression values (microarray intensities) of the *STK11-*mutated METABRIC samples with those of *STK11-*WT METABRIC samples in each of the three breast cancer types as the followings: 5 mutated vs. 634 WT for ER+/PR+, 4 mutated vs. 266 WT for ER+, and 4 mutated vs. 178 WT for TN. We used a group-comparing method available in the cBioPortal and detected 40 DEGs (6 up and 34 downregulated), 132 DEGs (26 up and 107 downregulated) and 139 DEGs (24 up and 115 downregulated) in ER+/PR+, ER+, and TN breast cancers, respectively (Source Data Fig. 3). The DEA revealed that these three breast cancer subtypes commonly have a higher portion of downregulated genes in *STK11-*mutated METABRIC samples when compared with *STK11-*WT METABRIC samples. The overall expression pattern of most DEGs was specific to each breast cancer subtype with limited overlap (Fig. 3d). For example, *HLA-DRB5* was the only common gene downregulated in *STK11*-mutated METABRIC samples in all three breast cancers (−1.47, −1.69 and −1.83 log_2_ fold changes in ER+/PR+, ER+ and TN, respectively) (Source Data Fig. 3). Furthermore, a GSEA identified a total of 40 GO terms enriched in the category of biological processes (16 in ER+/PR+, 14 in ER+, and 10 in TN) (Source Data Fig. 3). Interestingly, there were no common GO terms enriched among the three breast cancers, with the exception ‘Response to carbohydrate stimulus (GO:0009743)’ that was overrepresented in ER+/PR+ and ER+ breast cancers (Fig. 3e and Source Data Fig. 3). In addition, 30 of the 40 GO terms were enriched with downregulated genes (Fig. 3e) suggesting that mutations in *STK11* ultimately decrease functional activity of the diverse biological processes enriched in individual breast cancers.

**Fig. 3:**
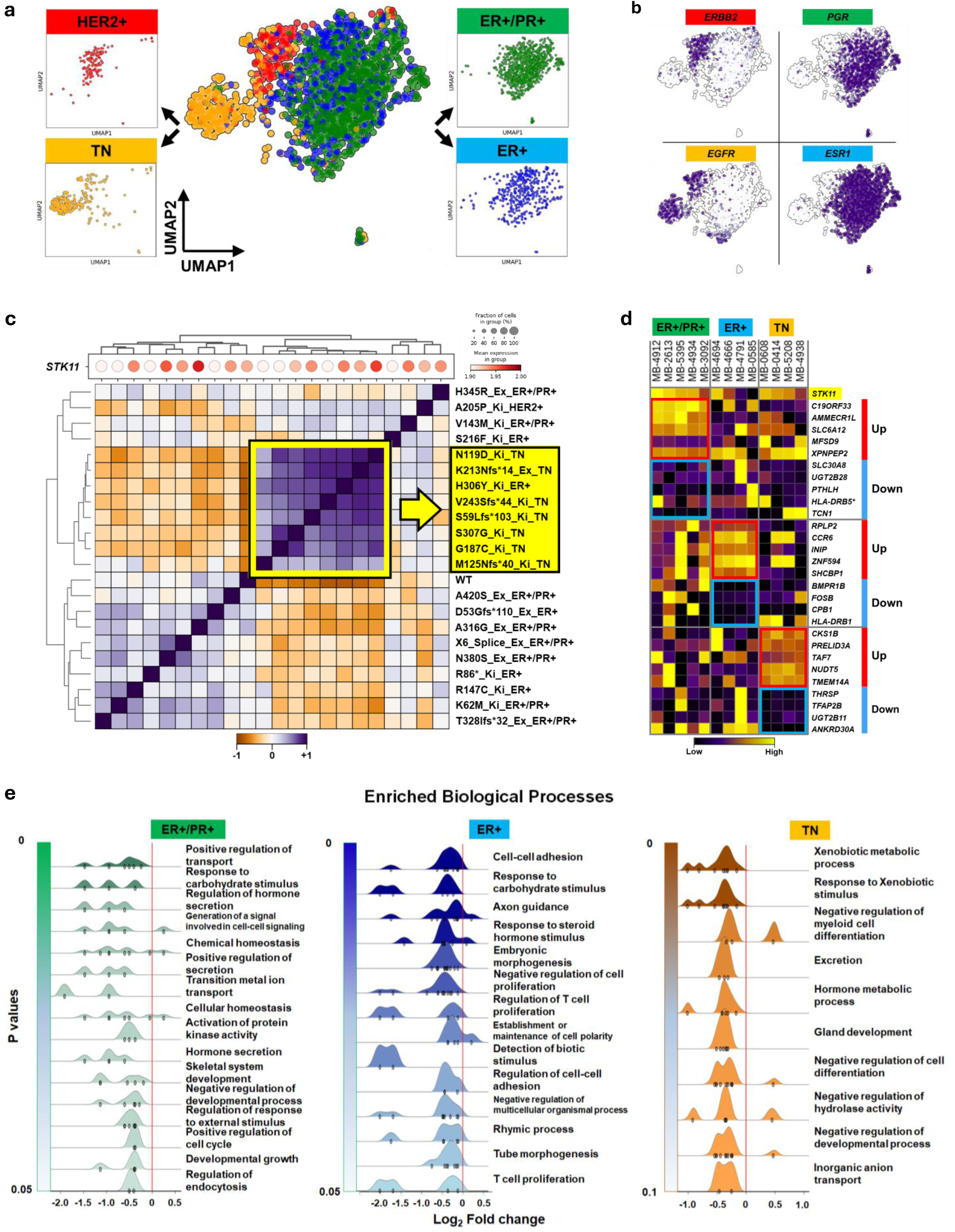
Profiles of differential gene expression at bulk-cell resolution in *STK11*-mutated breast cancer. **a**, Clustering analysis for breast cancers containing *STK11* mutation. Clusters are presented on a UMAP plot for METABRIC samples from four breast cancers, HER2+(red), ER+/PR+ (green), ER+ (blue) and TN (*BRCA1*-wild type triple negative, orange) that contain *STK11* mutations. Black arrows indicate individual breast cancers. **b**, Two HR and two RTK genes differentially expressed in the breast cancers in UMAP plots (*ESR1*: estrogen receptor 1, *PGR:* progesterone receptor, *ERBB2:* erb-b2 receptor tyrosine kinase 2, and *EGFR*: epidermal growth factor receptor, respectively). **c**, Correlation analysis of METABRIC samples. A white-red colour bar indicates low (1.90) and high (2.00) normalized mean expression of *STK11*, respectively. A gradient size of grey dots represents the fraction (%) of cells where *STK11* was expressed in individual METABRIC samples. Dendrogram trees and correlation matrix indicate hierarchical clusters of 1,980 METABRIC data that comprise 21 *STK11*-mutated and 1,959 *STK11*-wild type (WT) samples. The order of *STK11* mutation from top to bottom in the correlation matrix is symmetrical from left to right for the red-dot plot. An orange-white-purple colour bar represents low (−1.0), medium (0) and high (1.0) z-scaled correlation scores. The yellow-square line and arrow indicate highly correlated MEATBRIC samples containing eight *STK11* mutations, and the yellow arrow points to detailed information for seven mutations on kinase domain (Ki) and one mutation on exon (Ex) of *STK11* from TN and ER+ breast cancers. **d**, Differential gene expression in *STK11*-mutated breast cancers. Matrix plot presents normalized expression values (z-score of microarray intensities) for top 5 up or downregulated genes in *STK11*-mutated MEATBRIC samples when compared with *STK11*-WT METABRIC samples. Individual sample IDs per colour-matching breast cancer are presented above matrix plot, and gene symbols are listed at the right of the plot. Red and blue bars indicate up and downregulation of the genes, respectively. A purple-yellow colour bar represents low (purple) and high (yellow) z-scaled expression status of the genes, respectively. An asterisk (*) means that *HLA-DRB5* is one of the top 5 downregulated genes in ER+ and TN breast cancers as in ER+/PR+ breast cancer. **e**, Gene set enrichment analysis. Gene ontology terms in the category of biological processes that were overrepresented with DEGs in *STK11*-mutated MEATBRIC samples, when compared with *STK11*-WT METABRIC samples, are shown at the right of individual ridgeline plots for three breast cancers, ER+/PR+ (green), ER+ (blue) and TN (orange). X axis presents log_2_ fold change values of genes associated with enriched biological processes, and a gradient colour-matching bars in y axes indicate p-values at <0.05 for ER+/PR+ and ER+ breast cancers, and <0.1 for TN breast cancer. Red lines on ridgeline plots points to zero log_2_ fold change, that is, non-significant change. A height of individual ridgelines represents the number of genes involved in individual enriched biological processes.

### Detection of candidate biomarker genes based on association between mutational status of *STK11* and gene expression profiles in breast cancers

Overall, our results show that *STK11* mutations can induce a subtype-specific transcriptomic change amongst breast cancers. Here, we profiled an expression pattern of positively or negatively correlated genes with *STK11* expression to detect candidate biomarkers and characterize transcriptomes of breast cancers, particularly when *STK11* is mutated. We exemplified 20 genes that show positive (10 genes) and negative (10 genes) correlation with *STK11* expression in S*TK11*-WT METABRIC samples for each of ER+/PR+, ER+ and TN breast cancers (Fig. 4a). Importantly, the correlation directions of the exemplified genes are inversed in *STK11*-mutated METABRIC samples indicating differential expression as well as inverse activity of associated biological processes. We performed a GSEA on each of 6 gene sets (3 positively correlated gene sets and 3 negatively correlated gene sets; 10 *STK11*-correlated genes per gene set) using GO database and identified GO terms associated with mutational status of *STK11* per breast cancer. For example, two biological processes, ‘negative regulation of apoptotic process (GO:0043066)’ and ‘negative regulation of programmed cell death (GO:0043069)’, were enriched with 6 (*PTGS2*, *PAK5*, *PAK2*, *RB1*, *SIRT1* and *HSP90B1*) of the 10 negatively *STK11*-correlated genes in *STK11*-WT METABRIC samples (positively correlated in *STK11*-mutated METABRIC samples) from ER+/PR+ breast cancer (Extended Data Fig. 2a and Source Data Extended Data Fig. 2). This indicates that both apoptotic and programmed cell death processes are positively correlated with *STK11* expression in *STK11*-WT ER+/PR+ breast cancer. The two GO terms were overrepresented with 3 (*PRKAA2*, *BRAF* and *RB1*) of the 10 positively *STK11*-correlated genes in *STK11*-WT METABRIC samples (negatively correlated in *STK11*-mutated METABRIC samples) from ER+ breast cancer (Extended Data Fig. 2b and Source Data Extended Data Fig. 2). These inverse GO-enrichment profiles revealed that the apoptotic and programmed cell death processes are differentially associated with *STK11* expression in HR+ breast cancers. In previous sections, we identified a downregulation of apoptosis-, autophage- and programmed cell death-medicated signalling pathways in BT474_LKO and MDA-MB-231_LKO when compared with their corresponding parent cell lines (Source Data Fig. 1). *AEN* encodes for the protein, apoptosis enhancing nuclease, that contributes to apoptosis and autophagy in a p53-dependent manner^41^. We observed the downregulation of *AEN* in three epithelial cell types (luminal, mammary and myoepithelial) of most breast cancer subtypes from breast cancer patients (Source Data Fig. 2). Our findings in the METABRIC samples, combined with our observation in *STK11*-knockout cell lines and scRNA-seq dataset of breast cancer patients, strongly support LKB1 differentially regulates apoptotic-mediated programmed cell death processes in breast cancers. Furthermore, another two GO term, ‘negative regulation of TOR signalling (GO:0032007)’ and ‘negative regulation of cell cycle (GO:0045786)’, were significantly enriched with 3 (*PRKAA2, SIRT1, PRKACA, RPS6KA15*) and 5 (*RB1*, *SIRT1*, *BRSK1*, *ATF2* and *RPS6KA2*) of the 10 positively and negatively *STK11*-correlated genes in *STK11*-WT METABRIC samples from ER+ and TN breast cancers, respectively (Extended Data Fig. 2b,c and Source Data Extended Data Fig. 2). This indicates that cell cycle process is also differentially correlated with *STK11* expression when *STK11* is wild type in the two different breast cancers. In a precious section, a GSEA revealed a downregulation of cell cycle-related signalling pathways in BT474_LKO and MDA-MB-231_LKO (Source Data Fig. 1). Collectively, these GSEA results suggest that loss of LKB1 function differentially contributes to enhanced cell proliferation through differential regulation of apoptotic pathways, particularly in TN breast cancer. Both Retinoblastoma 1 (*RB1*) and Sirtuin1 (*SIRT1*) are tumour suppressors and known to negatively regulate cell cycle, cell proliferation and metabolism^42–44^. We found that the expression of these genes was negatively correlated with *STK11* expression in *STK11*-WT METABRIC samples for ER+/PR+ and TN breast cancers but positively correlated in *STK11*-WT METABRIC samples for ER+ breast cancer (Fig. 4a). Further studies are required to experimentally demonstrate how tumourigenesis in breast cancers attributes to correlation directions among LKB1, RB1 and SIRT1, and which molecular interactions are associated with those tumour suppressors in breast cancers.

**Fig. 4:**
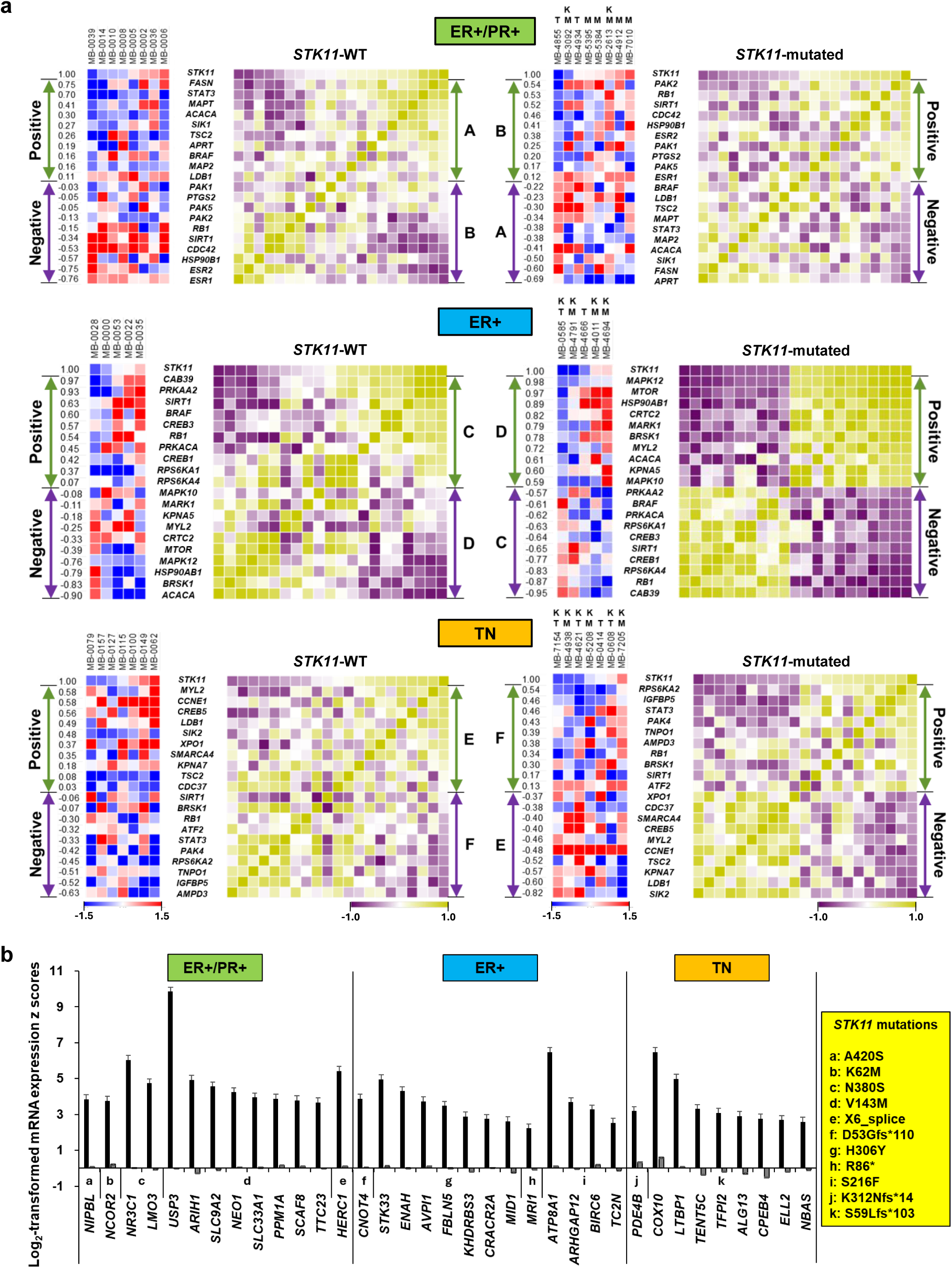
Genes associated with differential expression and *STK11* mutations in breast cancers. **a,** Candidate biomarker genes correlated with *STK11* expression. Matrix plots present correlation between *STK11* mutational status and 20 genes (blue-red colour combination), and within the genes against *STK11* expression (purple-yellow colour combination) in ER+/PR+, ER+ and TN (*BRCA1*-WT triple negative) breast cancers. The numbers at the left of blue-red matrix plots indicate Pearson correlation co-efficient values of individual genes against the mRNA expression level of *STK11* (K: *STK11* mutation on kinase domain, T: Truncating or non-sense mutation on *STK11*, and M: Missense mutation on *STK11*). Each of the 6 gene sets (A-F) contains 10 selected genes that are positively (green arrows) or negatively (purple arrows) correlated with *STK11* expression showing an inverse direction between *STK11*-WT and *STK11*-mutated METABRIC samples. Blue-red and purple-green colour bars indicate low (−1.5 for blue and −1.0 for purple) and high (1.5 for red and 1.0 for green) z-scaled correlation scores of individual genes, respectively. **b,** Candidate biomarker genes associated with *STK11* knockout. Individual pairs of black and grey bars indicate the highest and averaged mRNA expression z-scores of upregulated genes, respectively, in *STK11*-mutated METABRIC samples from ER+/PR+ (green), ER+ (blue) and TN (orange) breast cancers. X and y axes present candidate biomarker gene symbols and log_2_-scaled mRNA expression z-score values, respectively (See Source Data Fig. 5 for gene description). *STK11* mutations are listed up in the yellow text box. Standard errors for z-score values of individual genes are shown on the top of T-shape bars.

In addition to the introduced candidate biomarker genes showing the inverse-correlation directions associated with mutational status of *STK11* in the METABRIC samples, we also detected another set of 35 candidate biomarker genes that is directly associated with *STK11* knockout in mammary epithelial cells for breast cancers (Fig. 4b). These genes were selected from DEGs that were upregulated in the two *STK11*-knockout breast cancer cell lines, BT474_LKO and MDA-MB-231_LKO (Source Data Fig. 1). In more detail, we selected 675 commonly upregulated genes from the profiled DEGs in BT474_LKO and MDA-MB-231_LKO cells (Source Data Fig. 1), and used mRNA expression z-scores of the selected genes from the 13 *STK11*-mutated METABRIC samples of ER+/PR+, ER+ and TN breast cancers at early stage. We finally assigned 35 genes as candidate biomarkers of which mRNA expression z-score values were maximum in more than one of the METABRIC *STK11*-mutated samples when compared with those of METABRIC *STK11*-WT samples in each of the four breast cancers (Fig. 4b and Source Data Fig. 4). These mRNA-based biomarker genes can be clinically applicable for efficiently screening early-stage *STK11*-mutated breast cancers.

### Loss of LKB1 expression leads to mammary gland tumourigenesis

Lkb1 hypomorphic mice previously described^6^ were used to determine whether loss of LKB1 expression leads to mammary gland tumourigenesis in multiparous mice. In earlier studies using Lkb1*^fl/fl^*mice, expression of Lkb1 in mammary gland epithelial cells had not been reported^6^. Here, we confirm hypomorphic expression of Lkb1 in whole mammary gland tissue harvested from multiparous *Lkb1^+/+^, Lkb1^+/fl^, and Lkb1^fl/fl^* mice remained stable (Fig. 5a). To test whether loss of Lkb1 expression from mouse mammary glands resulted in tumourigenesis, we bred *Lkb1^fl/fl^*mice with whey acidic protein (WAP)-Cre transgenic mice (C57BL/6J) to give rise to *Lkb1^fl/fl^/*WAP-Cre. Mice were divided into two groups, those that remained nulliparous and those that were multiparous. The expression of the WAP-Cre transgene is largely limited to epithelial cells located at duct termini and within developing alveoli or luminal cells^28^. Since the WAP promoter is active during pregnancy and lactation, all long-term studies were conducted in multiple-birth females, unless otherwise stated. Kaplan-Meir curves (Fig. 5b) showing tumour-free survival indicate that by 11 months, 50% of multiparous *Lkb1^fl/fl^*/WAP-Cre mice developed tumours with 81% penetrance observed, compared with multiparous and nulliparous *Lkb1^fl/fl^* mice, despite being hypomorphic for Lkb1 expression, did not develop tumours over the duration of our study. Western blot analysis confirms loss of Lkb1 expression in multiparous *Lkb1^fl/fl^*/WAP-Cre mice, compared with expression in nulliparous Lkb1*^fl/fl^*/WAP-Cre mammary glands (Fig. 5c). Analysis of mammary glands from nulliparous *Lkb1^fl/fl^* and tumours from multiparous *Lkb1^fl/fl^*/WAP-Cre mice by IHC confirmed loss of Lkb1 expression. At 200 and 330 days, loss of Lkb1 expression is associated with ductal carcinoma *in situ* (middle panel), and invasive ductal carcinoma (bottom panel), respectively (Fig. 5d). To determine whether our *Lkb1^fl/fl^/*WAP-Cre mouse model presented with similar metabolic defects as our previous *Lkb1^−/−^/*NIC model^4, 5^, we conducted IHC analysis on mammary tissue from nulliparous and tumours from multiparous mice (Fig. 5e). IHC for expression of known targets of LKB1 including cyclin D1^8^, and c-Myc^3^ were elevated, as were phosphorylation of ribosomal protein S6 (pS6), a target of mTOR activation^45^, and metabolic markers, hypoxia inducible factor 1 (HIF1α)^46^, and lactate dehydrogenase (LDH)^4, 5, 47^. Furthermore, both ERα, a known binding partner of LKB1^3^ and PR were reduced, while ErbB2^4, 5^ expression was reduced and diffuse (Fig. 5e). In addition to characterizing Lkb1-known target gene expression in hypomorphic mice, we profiled expression patterns of those target genes in the two *STK11*-knockout cell lines comparing with their parent cells, and in the three epithelial cell types of breast cancer patients although the mutational status of *STK11* is unknown (Extended Data Fig. 3). As previously described, *PGR* and *ERBB2* showed upregulation in BT474_LKO, and *EGFR* was upregulated in MDA-MB-231_LKO and PGR (see Fig. 1e for HR and RTK genes in more detail). Interestingly, most target genes presented a similar expression pattern in *STK11*-knockout cells of the two cell lines when compared with their parent cells, except *LDHA* that was downregulated (−0.67 log_2_ fold change) in BT474_LKO but upregulated (0.21 log_2_ fold change) in MDA-MB-231_LKO (Source Data Fig. 1 and Extended Data Fig. 3a). Furthermore, the *LDHA* was also upregulated in luminal (0.48 log_2_ fold change) and mammary epithelial cells (1.25 log_2_ fold change) of TN breast cancer when compared with the two normal-corresponding epithelial cells (Source Data Fig. 2 and Extended Data. Fig. 3b).

**Fig. 5:**
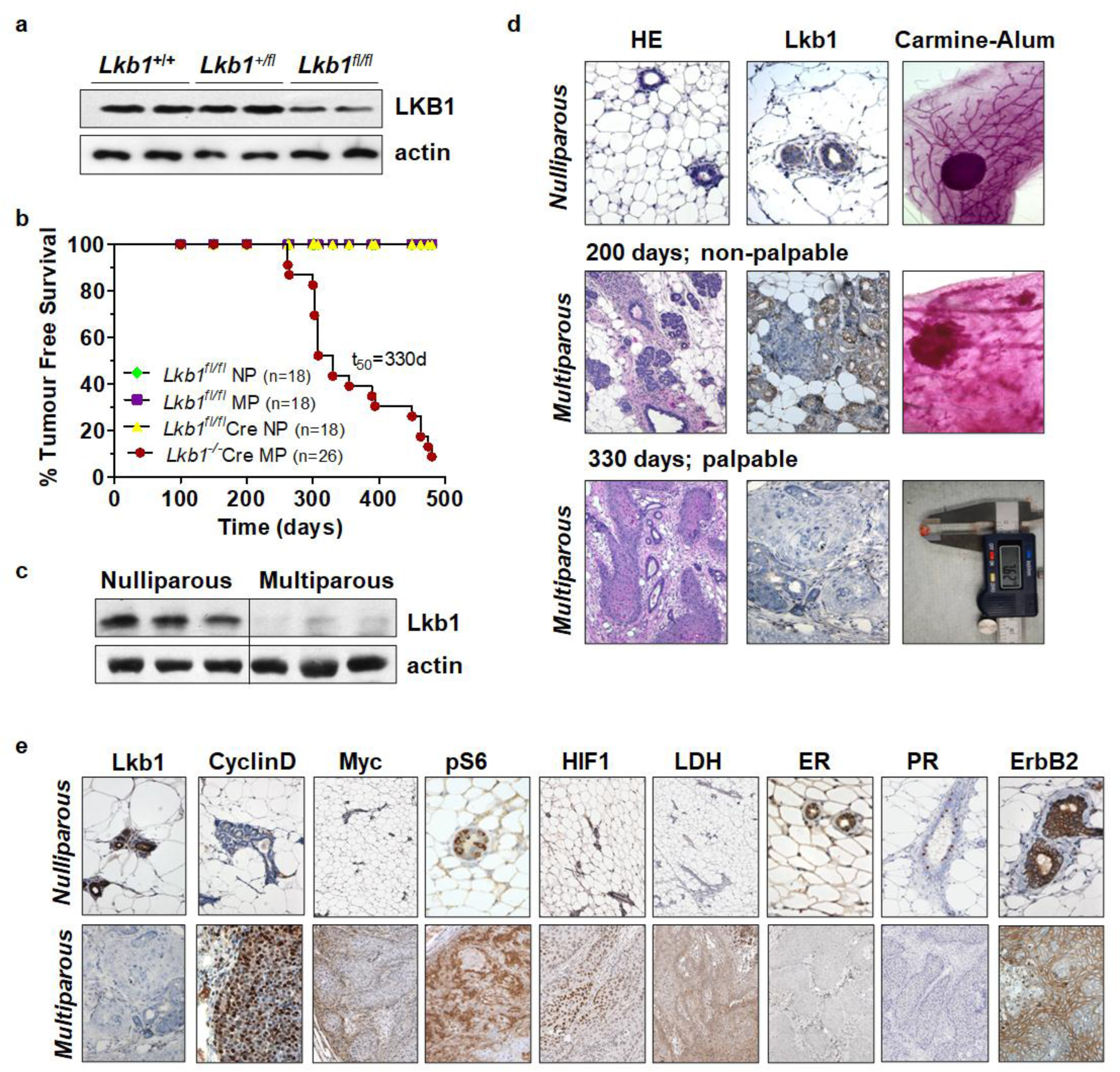
Loss of LKB1 expression results in mammary gland tumourigenesis. **a,** Expression of Lkb1 in hypomorphic mice. Mammary gland #4 was harvested from *Lkb1^+/+^, Lkb1^+/fl^*, and *Lkb1^fl/fl^*, mice and prepared for Western blot analysis for protein expression of LKB1, and actin (loading control) using anti-LKB1 and -actin antibodies. Two litter-mates are shown to highlight variation between mice. **b,** Kaplain-Meir percent tumour free survival curves for *Lkb1^fl/fl^*nulliparous (NP), *Lkb1^fl/fl^* multiparous (MP), *Lkb1^fl/fl^*,*/Cre* NP, and *Lkb1^−/−^/Cre* MP mice. Log Ranked test, p<0.0001. T_50_ represents time in days where 50% of the females presented with tumours and n is the number of female mice analysed. **c,** western blot analysis for Lkb1 expression in nulliparous and multiparous *Lkb1^fl/fl^*,*/Cre* NP, and *Lkb1^−/−^/Cre* MP mice, respectively. Data is representative of three separate mice to highlight variation between mice. **d,** IHC of Lkb1 expression in tumours from nulliparous and multiparous mice; H/E (10x), Lkb1 (10x), counter stained with haematoxylin. Wholemount Carmine alum staining, and representative calliper measurement highlighted. For multiparous mice, two time points indicated: 200 days non-palpable tumours and 330 days palpable tumours. **e,** Lkb1 (10x), Cyclin D1 (20x), c-Myc (10x), pS6 (40x), HIFa (10x), LDH (10x), ERα (20x), PR (10x), ErbB2 (40x). Counter stained with haematoxylin. Data is representative of six separate mice.

Overall, our *Lkb1^fl/fl^/*WAP-Cre model confirms that loss of *Lkb1* expression is adequate to drive mammary gland tumourigenesis with enhanced mTOR activity and cancer metabolism. Overall, these findings support results from our METABRIC analysis of *STK11* mutation status (Fig. 3) that suggests loss of LKB1 function differentially contributes to enhanced cell proliferation through differential regulation of signalling pathways, particularly in TN breast cancer.

## Discussion

The tumour suppressor LKB1 has long been implicated in diverse cancers through different pathways that include cellular energy metabolism, chromatin remodelling and polarity, just to name a few. Here, we characterize the consequences of LKB1 loss in breast cancer by integrating CRISPR-Cas9-mediated knockout in human cell lines, scRNA-seq data of breast cancer patients, bulk-tumour genomics analysis of the METABRIC cohort and a novel murine model. Together, the outcome of our research findings extends the biological scope of LKB1 well beyond its canonical AMPK-mTOR axis and establish context-specific transcriptomic programmes across breast cancer subtypes.

In human breast cancer cell lines, scRNAseq analysis of *STK11*-knockout BT474 (ER+/PR+/HER2+) and MDA-MB-231 (ER-/PR-/HER2−) cells revealed strikingly divergent transcriptomic profiles in response to LKB1 loss. Notably, loss of *STK11* enhanced the expression of *PGR* and *ERBB2* in BT474_LKO with non-significant change in MDA-MB-231_LKO, while *EGFR* expression showed inverse pattern between the two *STK*11-knockout cell lines. This suggests that LKB1 may function to regulate receptor tyrosine kinases and hormone receptor signalling. This is particularly intriguing in the context of therapeutic resistance, as amplified EGFR and HER2+ signalling are established drivers of endocrine and HER2−directed therapy failure^48^. In BTB474_LKO cells, we observed suppressed *FOS* expression and enhanced *PGR*, confirming an established role for FOS as a negative regulator of progesterone-mediated transcription^32^, thus positioning LKB1 as a positive upstream regulator of this signalling axis. Gene set enrichment analysis further illustrates the signalling consequences of LKB1 loss. The suppression of estrogen-dependent signalling pathways in both *STK11*-knockout cell lines, combined with significant downregulation of *GNAI1* in MDA-MB-231_LKO, a key mediator of G protein-coupled estrogen receptor signalling, supports our earlier findings where we identified LKB1 as a regulator on non-genomic estrogen signalling^3^. Conversely, androgen-mediated signalling pathways were downregulated following *STK11* knockout in both cell lines, with *BMPR1B* showing the most pronounced induction in BT474_LKO compared with MDA-MD-231_LKO. Given that AR signalling has recently emerged as a therapeutic target in TNBC^49, 50^, the relationship between LKB1 and this pathway may have important implications for patient stratification. In addition to *FOS*, among the most functionally relevant downregulated genes in response to *STK11* knockout were *IL32* and *IRF9* which are associated with the pathway of cytokine signalling in the immune system, suggesting a shared regulatory node between LKB1 tumour suppressor function and proinflammatory signalling. Co-expression analysis identified 92 *STK11*-coexpression genes, of which 17 are interconnected in a protein-protein interaction network associated with energy-dependent mTOR regulation through the LKB1-AMPK axis. Among these, *RHEB*, a direct activator of mTORC1, was among the top co-expressed genes, further underscoring a central role for LKB1 signalling in mTOR regulation. Importantly, *EIF4A2*, a mRNA helicase, associated with cell proliferation and chemosensitivity in TNBC^51^, was the sole *STK11*-coexpression gene upregulated in response to *STK11* knockout, consistent it the tumour suppressor function of LKB1.

Profiling of scRNAseq data from the four breast cancer patient subtypes, such as ER+, HER2+, TN (*BRCA1*-wild type) and TN-BR (*BRCA1*-mutated) revealed that tITH was highest in luminal epithelial cells and lowest in fibroblasts, with epithelial subpopulations showing the strongest correlation with *STK11* co-expression gene profiles (Fig. 3a). Important to note, is that TNBC displayed the most distinct pattern where *STK11* was markedly downregulated in mammary epithelial cells yet paradoxically, upregulated in myoepithelial cells. This cell-type selective LKB1 function resonates with emerging evidence that LKB1 exerts context-dependent roles in different epithelial lineages and suggests that myoepithelial upregulation of *STK11* expression in TNBC may represent a compensatory or reactive response to tumour microenvironments^52^.

Analysis of 1,980 METABIC samples confirmed that *STK11* mutations are enriched in TNBC and disproportionately affect the catalytic domain, with 85.7% of TN-associated mutations localized with this region. This distribution aligns with the expectation that catalytic deficient mutants^8^, which retain the ability to scaffold signalling complexes but lack kinase activity, may exert dominant-negative or gain-of-function effects distinct from simple loss of expression^3, 53–55^. The single shared DEG across ER+, ER+/PR+, and TN *STK11*-mutated samples was *HLA-DRB5*, whose consistent downregulation across breast cancer subtypes may reflect LKB1-dependent immune surveillance mechanisms. Loss of MHC class II antigen presents an established mechanism for immune evasion in breast cancer^56, 57^, and these findings suggest that catalytic deficient mutants of LKB1 may contribute to an immunosuppressed tumour microenvironment.

Our *Lkb1^fl/fl^*/WAP-Cre mouse model provides evidence *in vivo* that the loss of Lkb1 is adequate and sufficient to drive mammary gland tumourigenesis with a long latency in multiparous mice. Previous work by Xu et al, found that conditional knockout of *Brca1* exon 11 in murine mammary epithelial cells resulted in tumour formation with long latency between 10-13 months^58^, whereas Ludwig et al, found Cre-mediated deletion of *Brca2* exon 3 and 4 in murine mammary epithelial cells resulted in tumours with a long latency of approximately 1.4 years^59^. In our model, we observe elevated mTOR signalling, and enhanced metabolism, recapitulating the hypermetabolic phenotype we previously described in our *Lkb1^−/−^/*NIC murine model of breast cancer^4, 5^. This metabolic reprogramming is consistent with established role for LKB1 in regulating mTOR activity through AMPK^4, 5, 45^. The enhanced expression of c-Myc and cyclin D1 in our LKB1-deficient tumours further supports prior mechanistic studies linking LKB1 catalytic activity to oncogenic transcriptional regulation that destabilizes epithelial integrity^3, 8, 60^. Furthermore, the reduced expression of ERα, PR and ErbB2 in this model suggests loss of *Lkb1* expression is likely to play a role in the development and/or progression of TNBC, consistent with our finding from analysis of *STK11*-mutated METABRIC.

We applied an integrated approach to transcriptomic characterization of *STK11* expression and activity from multiple biological breast cancer sources and showed that loss of *Stk11* expression is sufficient to drive mammary gland tumourigenesis in a murine model. Profiling *STK11*-coexpression genes in breast cancer patients reveals that individual cell types differentially responded to transcriptomic changes in breast cancer, particularly in epithelial cells from TNBC. The uniqueness of gene expression pattern in TNBC is also demonstrated by identifying relationships between gene expression profiles and mutation status of *STK11* at bulk-cell level. The 35 candidate biomarker genes identified from the intersection of *STK11*-knockout transcriptomic data and *STK11*-mutated METABRIC samples represent clinically actionable gene panel for early-detection screening of *STK11-*altered breast cancers at mRNA resolution. We conclude that LKB1 is associated with more diverse biological processes than those reported from previous cancer studies.

## Supporting information

Source Data Fig. 1. scRNA-seq from STK11-knockout cells

Source Data Extended Fig. 1. GSEA for STK11-knockout cells

Source Data Fig. 2. scRNA-seq from breast cancer patients

Source Data Extended Fig. 2. GSEA for METABRIC Marker

Source Data Fig. 3. Microarray from METABRIC samples

Source Data Fig. 4. Candidate biomarker genes

## Acknowledgements

We thank former members of the Marignani Discovery Research Laboratory for their support. This work was supported by CIHR (MOP-67039), Nova Scotia Health Research Foundation, Breast Cancer Canada and by private donations administered through the Dalhousie Medical Research Foundation. The authors acknowledge that Dalhousie University sits on Mi’kma’ki, the ancestral and unceded territory of the Mi’kmaq People. We are all Treaty People. We recognize that African Nova Scotians are a distinct people whose histories, legacies and contributions have enriched that part of Mi’kma’ki known as Nova Scotia for over 400 years.

## Supplement table legend

**Supplement table 1. Information on scRNA-seq NGS data of human breast cancer cell lines.** The table contains scRNA-seq NGS data generated from four human breast cancer cell lines, BT474, *STK11*-knockout BT474 (BT474_LKO), MDA-MB-231 and *STK11*-knockout MDA-MB-231 (MDA-MB-231_LKO).

## Extended data figure legends

**Extended Data Fig. 1.**
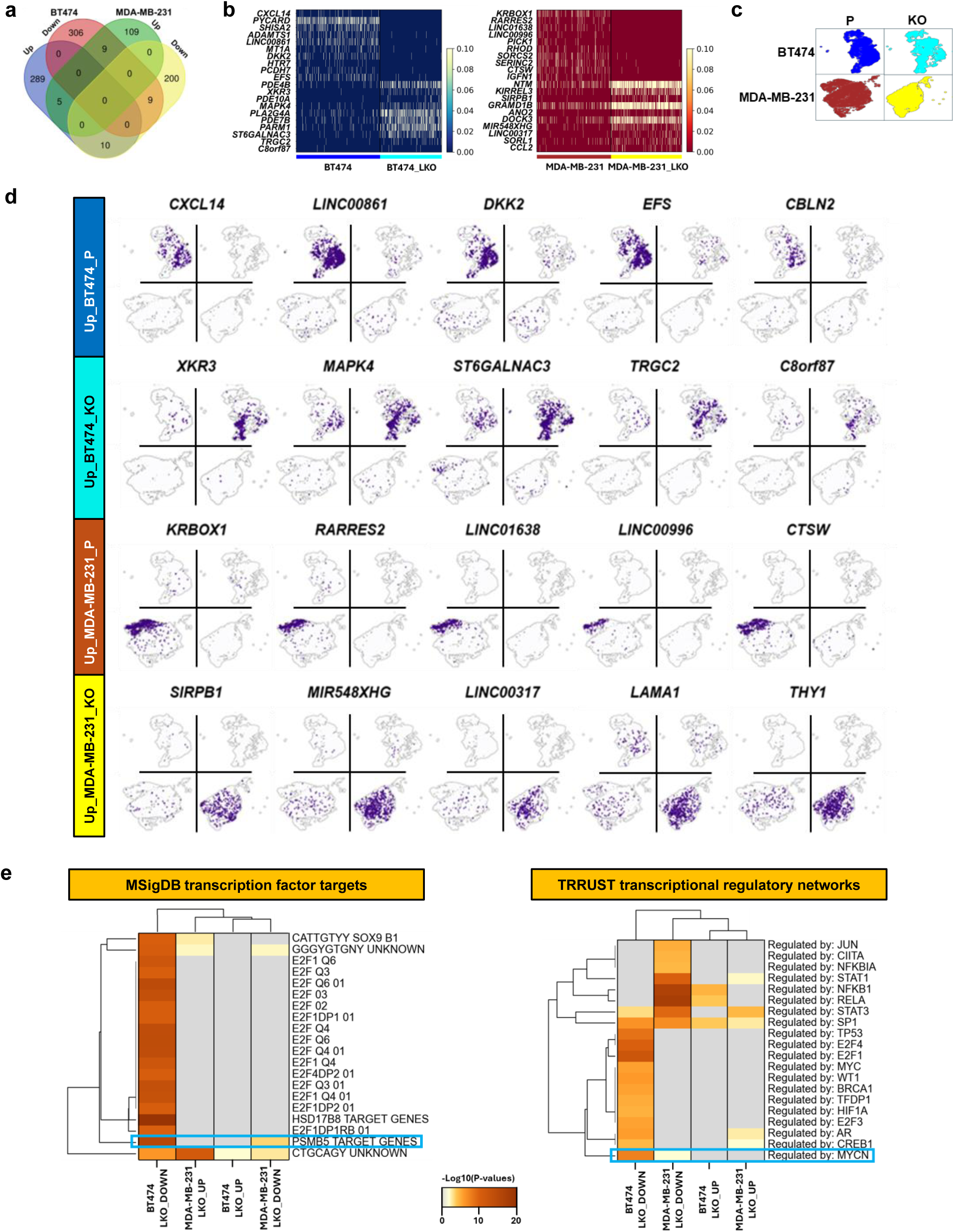
Profiles of gene expression in *STK11*-knockout cells compared with parent cells for each of BT474 and MDA-MB-231 breast cancer cell lines. **a,** Differentially expressed genes in *STK11*-knockout cells. Numbers in individual colour sections of Venn diagram indicate the number of genes up (≥ 1 log_2_ fold change) or downregulated (≤ −1 log_2_ fold change) in *STK11*-knockout cells compared with parent cells. **b**, The left of matrix plots shows the list of the most significantly upregulated top 10 genes in *STK11*-knockout cells of BT474 (BT474_LKO; left) and MDA-MB-231 (MDA-MB-231_LKO; right) when compared with each of their corresponding parent cell lines. The blue-green-yellow and brown-orange-yellow colour bars represent low (0 for blue and brown) and high (0.10 for yellows) z-scaled normalized expression values of the genes, respectively. **c**, Distribution of single cells from four breast cancer cell lines. Individual UMAP plots indicate BT474 (blue), BT474_LKO (cyan), MDA-MB-231(brown) and MDA-MB-231_LKO (yellow) cell lines. P and KO represent parent cells and *STK11*-knockout cells, respectively. **d**, Individual UMAP plots present single cells where 5 genes per cell line are cell line-specifically expressed (*CXCL14*: C-X-C motif chemokine ligand 14, *LINC00861*: long intergenic non-protein coding RNA 861, *DKK2*: dickkopf WNT signaling pathway inhibitor 2, *EFS*: embryonal Fyn-associated substrate, *CBLN2*: cerebellin 2 precursor, *XKR3*: XK related 3, *MAPK4*: mitogen-activated protein kinase 4, *ST6GALNAC3*: ST6 N-acetylgalactosaminide alpha-2,6-sialyltransferase 3, *TRGC2*: T cell receptor gamma constant 2, *C8orf87* (*LINC02906*): long intergenic non-protein coding RNA 2906, *KRBOX1*: KRAB box domain containing 1, *RARRES2*: retinoic acid receptor responder 2, *LINC01638*: long intergenic non-protein coding RNA 1638, *LINC00996*: long intergenic non-protein coding RNA 996, *CTSW*: cathepsin W, *SIRPB1*: signal regulatory protein beta 1, *MIR548XHG*: MIR548X host gene, *LINC000317*: long intergenic non-protein coding RNA 317, *LAMA1*: laminin subunit alpha 1, *THY1*: Thy-1 cell surface antigen). **e**, Enriched transcription factors in four breast cancer cell lines. Left and right heatmap plots show transcription factors enriched by GSEAs using MSigDB TFT and TRRUST database, respectively. Up or downregulated gene sets in the two *STK11*-knockout cell lines were used as input for the GSEAs and presented at the bottom of the plots. Grey-brown colour bar indicates negative log_10_(p-values) at the low (0), medium (10) and high (20) extent of statistical significance.

**Extended Data Fig. 2.**
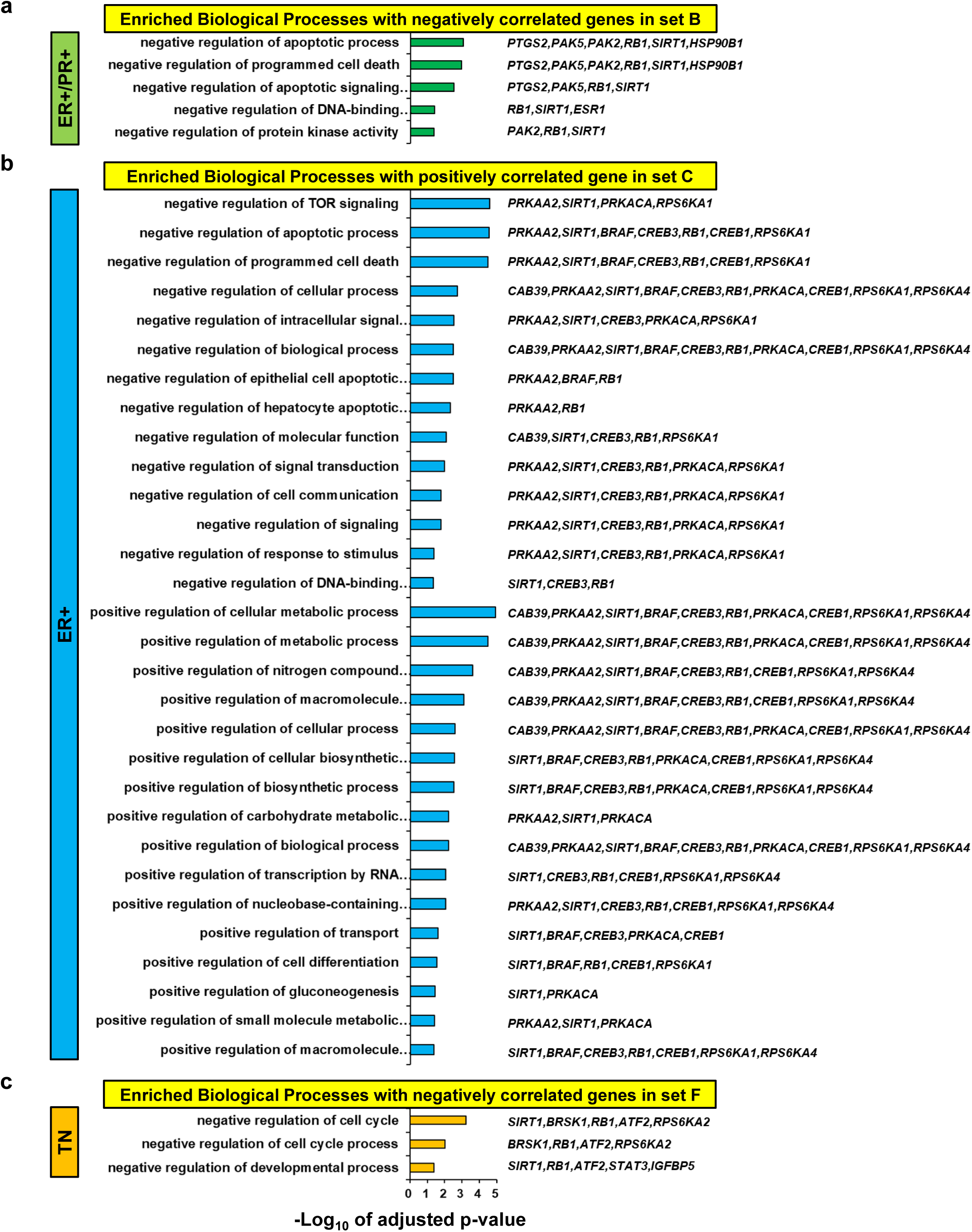
Gene set enrichment analysis with *STK11*-correlated genes in breast cancers. **a-c,** Biological processes positively or negatively regulated in ER+/PR+ (green; **a**), ER+ (blue; **b**) and *BRCA1*-wild type (WT) triple negative (TN, orange; **c**) breast cancers. Gene set B indicates genes negatively correlated with *STK11* expression in *STK11*-WT METABRIC samples from ER+/PR+ breast cancer, while Gene set C and F include genes positively correlated with *STK11* expression in *STK11*-WT METABRIC samples for ER+ and TN breast cancers, respectively (Fig. 5a). Colour-matching bars present -log_10_-adjusted p-values. Enriched GO terms and associated genes are presented left and right of the bar plot, respectively (see Source Data Extended Data Fig. 2 for full description of GO terms and genes).

**Extended Data Fig. 3.**
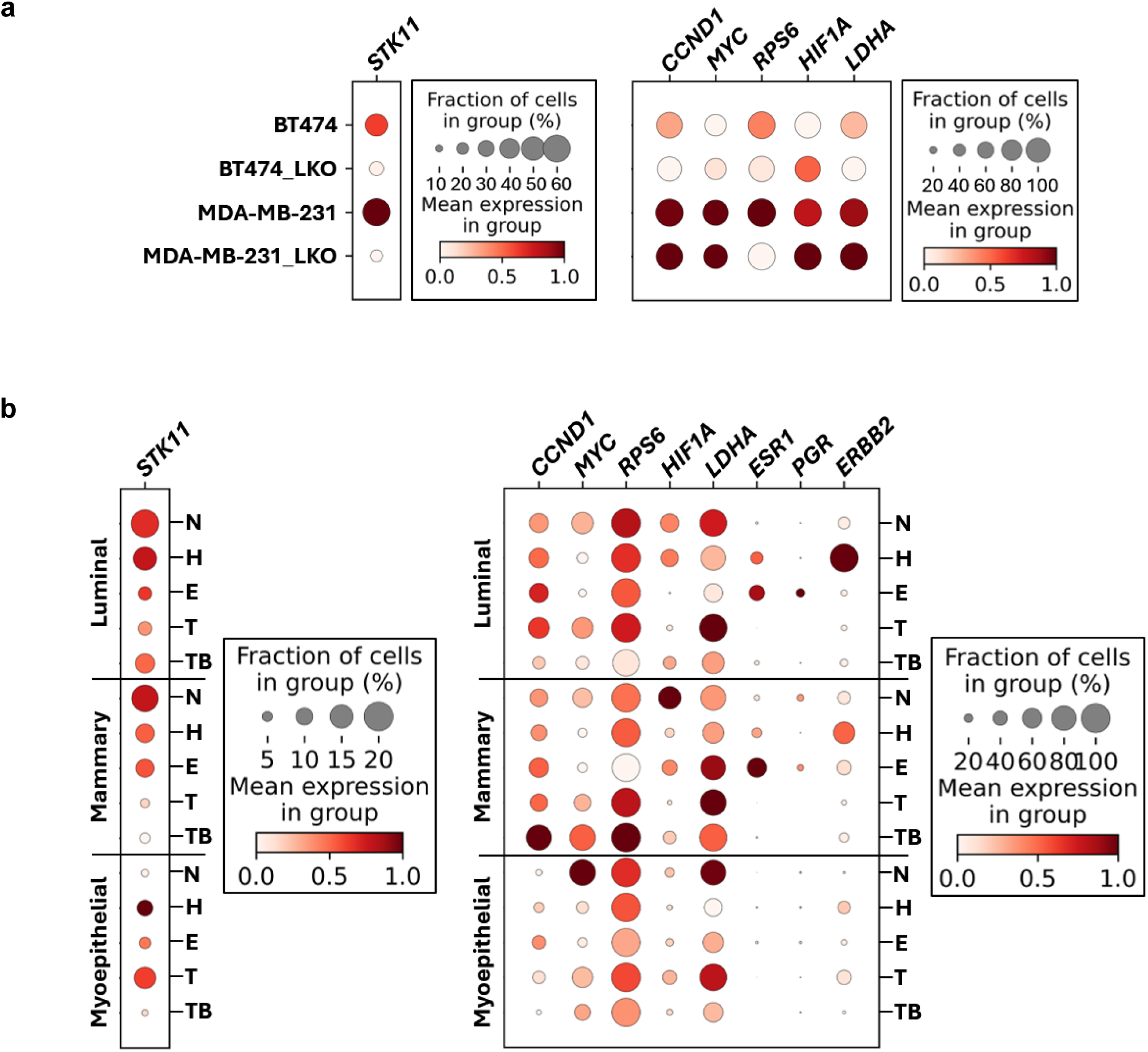
Differential expression profiles of functionally *STK11*-associated genes. **a-b**, Panels right to dot plots indicate a fraction (%) of cells according to a gradient size of dots for 6 genes in each of the four cell lines (BT474, BT474_LKO, MDA-MB-231, and MDA-MB-231_LKO) (Extended Data Fig. 3a) and 9 genes in luminal epithelial (top), mammary epithelial (middle) and myoepithelial (bottom) cell types from the normal mammary (N) and four different breast cancers (ER+: E, HER2+: H, *BRCA1*-wild type TN: T, and *BRCA1*-mutated TN: TB) (Extended Data Fig. 3b). The panels also represent a variety of mean expression values by a white-dark brown colour bar for low (0), middle (0.5) and high (1.0). See the Fig. 1e for an expression pattern of HR and RTK genes in the four cell lines (*STK11*: serine/threonine kinase 11, *CCND1*: cyclin D1, *MYC*: MYC proto-oncogene, bHLH transcription factor, *RPS6*: ribosomal protein S6, *HIF1A*: hypoxia inducible factor 1 subunit alpha, *LDHA*: lactate dehydrogenase A, *ESR1*: estrogen receptor 1, *PGR*: progesterone receptor, *ERBB2*: erb-b2 receptor tyrosine kinase 2).

## Notes

The authors declare no conflict of interest.

### Competing Interest Statement

The authors have declared no competing interest.

